# An Effective CTL Peptide Vaccine for Ebola Zaire Based on Survivors’ CD8+ Targeting of a Particular Nucleocapsid Protein Epitope with Potential Implications for COVID-19 Vaccine Design

**DOI:** 10.1101/2020.02.25.963546

**Authors:** CV Herst, S Burkholz, J Sidney, A Sette, PE Harris, S Massey, T Brasel, E Cunha-Neto, DS Rosa, WCH Chao, R Carback, T Hodge, L Wang, S Ciotlos, P Lloyd, R Rubsamen

## Abstract

The 2013-2016 West Africa EBOV epidemic was the biggest EBOV outbreak to date. An analysis of virus-specific CD8+ T-cell immunity in 30 survivors showed that 26 of those individuals had a CD8+ response to at least one EBOV protein. The dominant response (25/26 subjects) was specific to the EBOV nucleocapsid protein (NP). It has been suggested that epitopes on the EBOV NP could form an important part of an effective T-cell vaccine for Ebola Zaire. We show that a 9-amino-acid peptide NP44-52 (YQVNNLEEI) located in a conserved region of EBOV NP provides protection against morbidity and mortality after mouse adapted EBOV challenge. A single vaccination in a C57BL/6 mouse using an adjuvanted microsphere peptide vaccine formulation containing NP44-52 is enough to confer immunity in mice. Our work suggests that a peptide vaccine based on CD8+ T-cell immunity in EBOV survivors is conceptually sound and feasible. Nucleocapsid proteins within SARS-CoV-2 contain multiple class I epitopes with predicted HLA restrictions consistent with broad population coverage. A similar approach to a CTL vaccine design may be possible for that virus.

## 1. Introduction

Development of safe and effective vaccines for some viruses such as HIV and EBOV has been challenging [19]. Although vaccine development has been almost exclusively focused on eliciting a humoral immune response in the host through inoculation with whole protein antigen [51][69][59][29], CTL peptide vaccines producing a T-cell response may offer an important alternative approach [23]. For HIV and EBOV and influenza in particular, the potential of CTL vaccines has been discussed [21][7][56]. Although computational prediction alone has been used for T-cell vaccine design [2][14], we saw a unique opportunity to see if a preventative EBOV T-cell vaccine could be successfully designed based on the specific epitopes targeted by survivors of documented EBOV infection.

The notion of HLA restricted HIV control has been described [58]. Pereyra-Heckerman conducted an analysis of virus-specific CD8+ T-cell immunity in individuals living with HIV [43]. They reported that HIV controllers, individuals living with HIV not undergoing treatment who do not progress to AIDS, have CD8+ cells targeting different HLA restricted class I epitopes on HIV compared with progressors, individuals with HIV who progress to AIDS in the absence of therapy. Pereyra-Heckerman suggested that this observation could guide the *in-silico* development of a CTL vaccine for HIV and other diseases.

Acquired immunity has been documented after EBOV infection [4]. Antibody as well as T-cell responses have been described [44]. Sakebe et al. have shown that of 30 subjects surviving the 2013-2016 EBOV outbreak in West Africa, CD8+ T-cells from 26 of those survivors responded to at least one EBOV antigen, with 25 of the 26 responders targeting epitopes on EBOV NP [50]. One of the most commonly targeted EBOV eptitopes on EBOV NP in the survivor group (targeted by CD8+ cells from four survivors) was NP41-60 (IPVYQVNNLEEICQLIIQAF). They also suggested that a CTL vaccine could be designed using epitopes targeted by CD8+ T-cells identified in these EBOV controllers.

Human pathogen-derived peptide antigens that are also recognized by C57BL/6 T-cells have been previously described. These include peptides from vesicular stomatitis virus (VSV) RGYVYQGL [68], and human immunodeficiency virus (HIV) RGPGRAFVTI [5]. The existence of such epitopes makes a range of pre-clinical vaccine experiments possible without having to rely on non-human primates and expensive and complex-to-manage humanized mouse models. Wilson et al. showed that the EBOV nucleoprotein (NP) is an immunogen that provides protective, CTL-mediated immunity against EBOV in a C57BL/6 mouse model and that this protection was conferred by a peptide sequence within Ebola Zaire: NP43-53 (VYQVNNLEEIC) [73]. Wilson et al. came to this conclusion based on studying splenocytes harvested from mice vaccinated with Ebola Zaire NP using a Venezuelan equine encephalitis (VEE) vector. Their experiments showed that splenocytes from the vaccinated mice re-stimulated with NP43-53 had high levels of cytotoxic activity against target cells loaded with the EBOV NP peptide. Remarkably, NP43-53 also happens to be an 11 amino acid subsequence of the epitope identified by Sakebe et al. as most commonly favored for T-cell attack by survivors of the 2013-2016 EBOV outbreak in West Africa.

We set out to see if we could drive CTL expansion directed against NP43-53 to occur after vaccinating C57BL/6 mice with Ebola Zaire NP43-53 (VYQVNNLEEIC), and to subsequently conduct an *in-vivo* EBOV challenge study to see if this peptide was protective.

We fabricated adjuvanted microspheres for this study as a room temperature stable dry powder using the Flow Focusing process to be 11 *μM* in diameter so as to prevent more than one microsphere from being phagocytosed by any given antigen presenting cell (APC) at the same time [37]. By loading only one peptide sequence per microsphere, we maximized the peptide payload and mitigated the possibility of multiple, different peptide sequences being delivered to the APC simultaneously, which could possibly result in competitive inhibition at the motif which could interfere with antigen presentation and subsequent T-cell expansion (Supplementary Material Section 1).

We also set out to see if a similar approach to a CTL vaccine design for SARS-CoV-2 would be feasible based on an analysis of the HLA binding characteristics of peptide sequences on SARS-CoV-2 nucleocapsid.

## 2. Results

We used a previously described biodegradable dry powder, PLGA microsphere, synthetic vaccine platform adjuvanted with TLR-4 and TLR-9 agonists for this study [48]. In that article, we showed that the TLR-4 and TLR-9 agonists given together with a peptide in a mouse model did not produce T-cell expansion by ELISPOT and that microencapsulation of the peptide and the TLR-9 ligand, with the TLR-4 ligand in the injectate solution, was required to elicit an immune response to the delivered peptide antigen as determined by ELISPOT. That study also demonstrated that the microencapsulated peptides alone were insufficient to induce an adequate immune response without the presence of the TLR-4 and TLR-9 agonists administered as described. The TLR agonists used for this vaccine formulation are used in FDA approved vaccines and can be sourced as non-GMP or GMP material for pre-clinical and clinical studies.

We show here that the H2-D^b^ restricted epitopes VSV (RGYVYQGL) and OVA (SIINFEKL), when administered to C57BL/6 mice, each produce a CD8+ ELISPOT response to the administered peptide antigen with no statistically significant CD4+ response measurable by ELISPOT as shown in Figure 2c, and Figure 2d.

**Figure 1:**
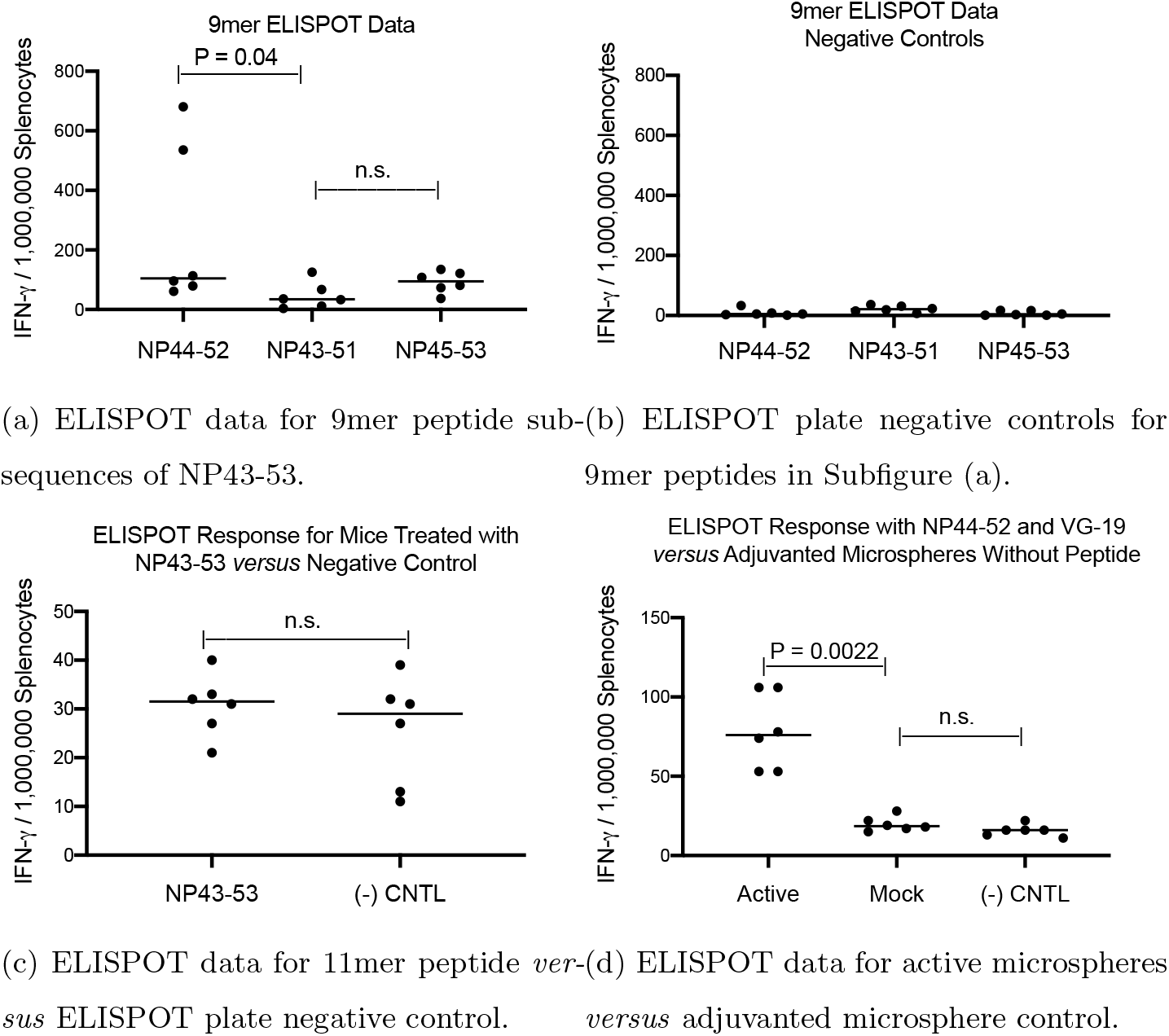
ELISPOT data from three groups of six mice each. Each of the three groups of mice were vaccinated (2mg adjuvanted microspheres via ID tail injection) with a different 9mer peptide sub-sequence of NP43-53. ELISPOT data showed NP44-52 produced the best immune response (1a). Mice vaccinated (2mg adjuvanted microspheres via ID tail injection) with the NP43-53 11mer produced the same immune response as ELISPOT plate negative control (1c). The same active formulation administered to mice for the challenge study (20mg adjuvanted microspheres via intraperitoneal injection) produced a positive immune response compared with both adjuvanted microsphere and ELISPOT plate controls (1d). (n.s. = not significant)

**Figure 2:**
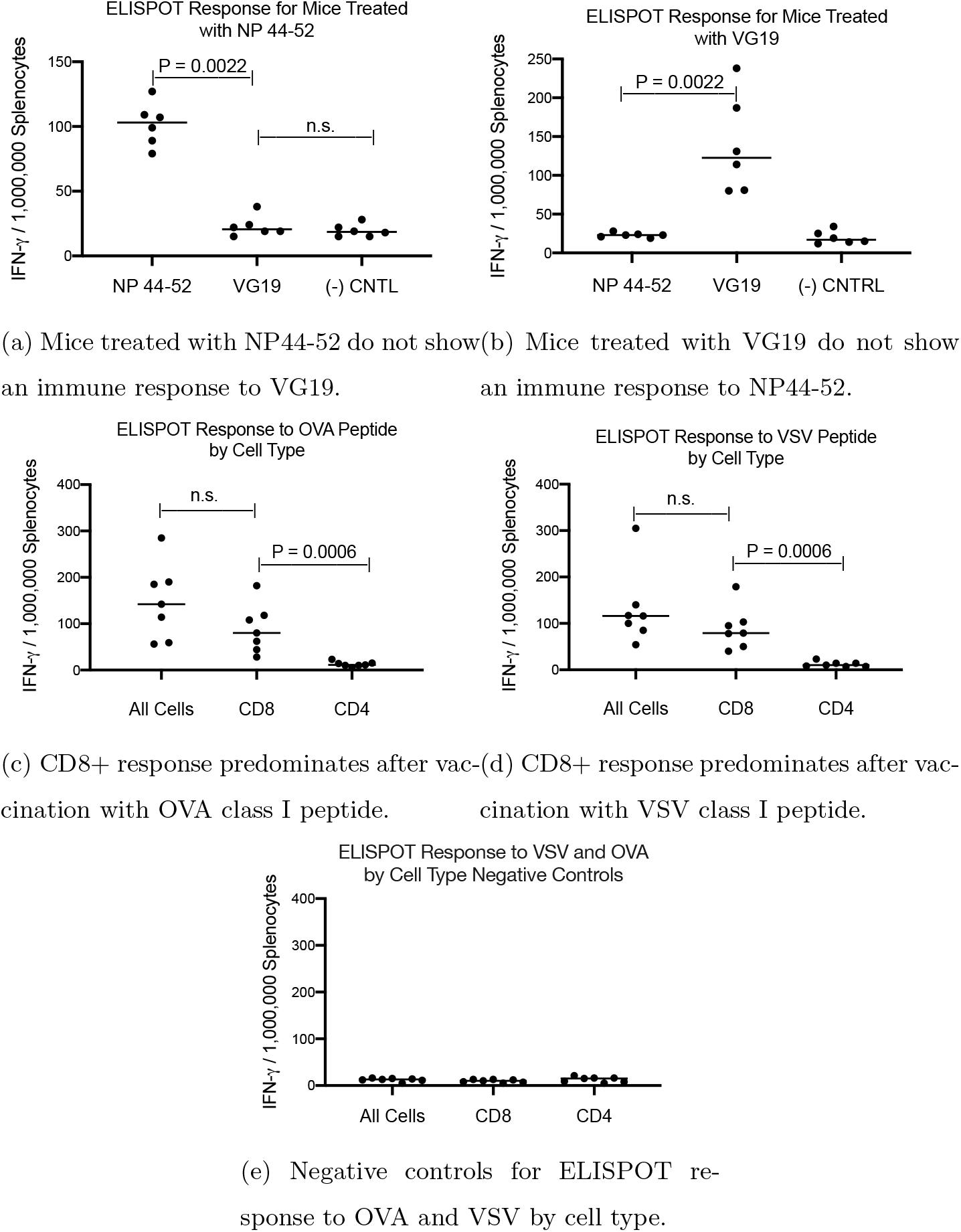
Six mice treated with NP44-52 (2mg adjuvanted microspheres via ID tail injection) were evaluated for their ELISPOT response to NP42-52 and VG19 (2a), and another group of six mice treated with VG19 (2mg adjuvanted microspheres via ID tail injection) have their ELISPOT responses to NP42-52 and VG19 shown in Figure 2b. In each of these groups, the mice generated an immune response only to the vaccinated peptide. A group of seven mice was evaluated for their immune response by cell type (using magnetic bead separation) for their ELISPOT responses evaluating total, CD8, and CD4 cell populations after vaccination (2mg adjuvanted microspheres via ID tail injection) with OVA peptide (2c) and VSV peptide (2d). For both peptides, the immune response by E1LISPOT was from the CD8+ cell population. (n.s. = not significant)

We used this adjuvanted microsphere peptide vaccine platform to immunize C57BL/6 mice with NP43-53, the CTL+ class I peptide antigen from the Ebola Ziare NP protein identified as protective by Wilson et al. [73]. Microspheres containing NP43-53 and CpG were prepared as a dry powder formulation and suspended before use in a PBS injectate solution containing MPLA, and administered intradermally via injection at the base of the tail into mice as described in a previous publication [48]. As illustrated in Figure 1c, there was no statistically significant difference between the ELISPOT data for the vaccinated mice *versus* the response seen in the negative ELISPOT controls.

Wilson reported that protection seen in her experiment was due to a peptide sequence within NP-43-53. We hypothesized that the NP43-53 epitope was inefficiently processed into MHC binding sub-sequences during antigen presentation. In order to explore possible H2-D^b^ matches for peptide sequences contained within Ebola Zaire NP43-53 (VYQVNNLEEIC), we prepared three peptide vaccine formulations, each containing one of the three possible 9mer sub-sequences within NP43-53. These sequences are shown in Table 1. We then vaccinated, via intradermal (tail) injection, three groups of mice with microspheres containing one of the three 9mer sub-sequences of NP43-53 (6 per group). ELISPOT analysis was performed, stimulating harvested splenocytes with the three possible 9mer sub-sequences. Splenocytes from mice receiving the NP44-52 sub-sequence had a statistically higher ELISPOT response than mice vaccinated with the other two possible sub-sequence 9mers (P < 0.0001) as shown in Figure 1a. This is consistent with the predicted H2-D^b^ binding affinity of YQVNNLEEI as shown in Supplementary Material Table 3.

**Table 1:**
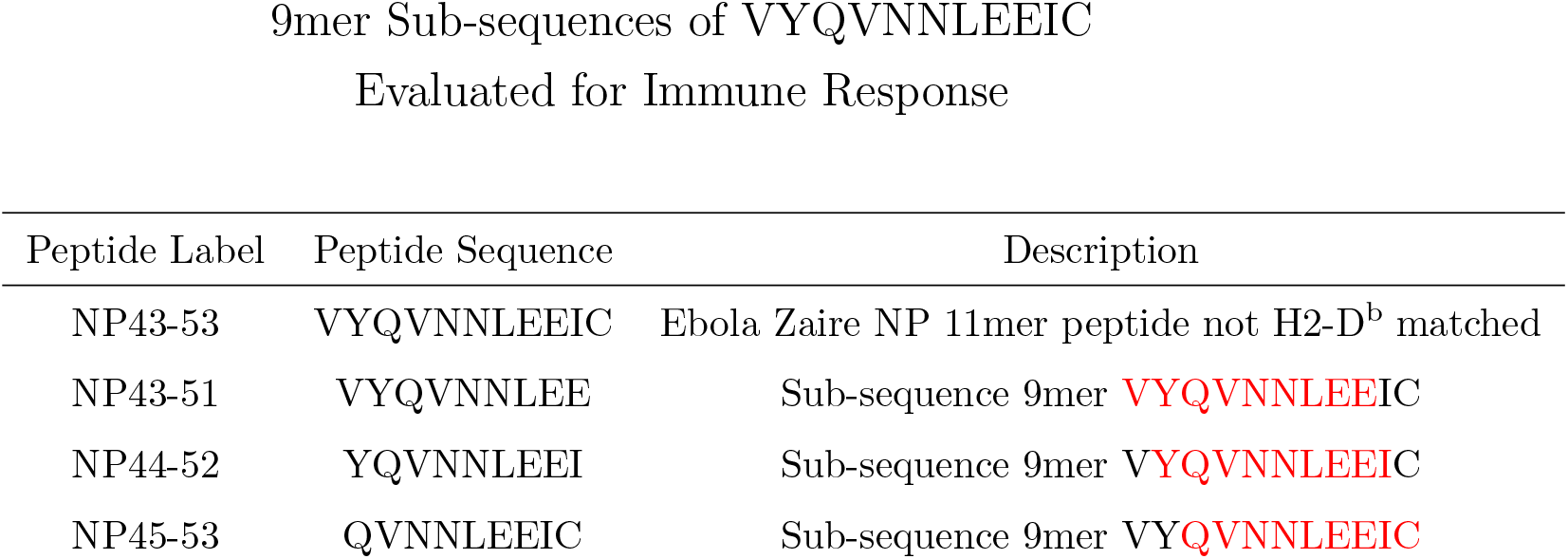
Class I peptides used in the study. NP43-53 is the class I 11mer described by Wilson et al. which we found not to produce an immune response in a C57BL/6 mouse model. NP43-51, NP 44-52 and NP 45-53 are the three possible 9mer sub-sequences of NP43-53.

| Peptide Label | Peptide Sequence | Description |
| --- | --- | --- |
| NP43-53 | VYQVNNLEEIC | Ebola Zaire NP 11mer peptide not H2-D<sup>b</sup> matched |
| NP43-51 | VYQVNNLEE | Sub-sequence 9mer **VYQVNNLEEIC** |
| NP44-52 | YQVNNLEEI | Sub-sequence 9mer **VYQVNNLEEIC** |
| NP45-53 | QVNNLEEIC | Sub-sequence 9mer **VYQVNNLEEIC** |

We then loaded one population of adjuvanted microspheres with NP44-52 and a second population of adjuvanted microspheres loaded with VG19 from EBOV Zaire NP 273-291 (VKNEVNSFKAALSSLAKHG), a Class II epitope predicted to be relevant to NP43-53 based on the TEPITOPE algorithm using a technique described by Cunha-Neto at al [14]. This peptide has a predicted favorable H2-I^b^ binding affinity as shown in Supplementary Material Table 5.

We showed that vaccination of 6 mice with the adjuvanted microsphere vaccine loaded with VG19 and NP44-52 showed an ELISPOT response to NP44-52 whereas 6 mice vaccinated with adjuvanted microspheres not loaded with peptide did not (Figure 1d).

We also showed that mice vaccinated with VG19 alone did not show an ELISPOT response to NP44-52 (Figure 2a) and, conversely, mice vaccinated with NP44-52 did not show a response to VG19 (Figure 2b).

We conducted a pilot study demonstrating that intraperitoneal injection of the adjuvanted microsphere vaccine produced a statistically superior immune response by ELISPOT compared with the same dose delivered by intradermal tail or intramuscular injection in C57BL/6 mice (Supplementary Material Section 2). Based on the data from that study, and the fact that the volume of the intraperitoneal space would allow larger amounts of microsphere suspension to be delivered, we chose to proceed with intraperitoneal administration for the challenge portion of this study delivering 20mg of microspheres per dose.

We dosed three groups of mice, ten mice per group, with the adjuvanted microsphere vaccine formulation containing NP44-52 and VG-19, with each peptide in a distinct microsphere population, and challenged these mice 14 days after vaccine administration with escalating IP administered doses of mouse adapted EBOV (maEBOV) (Group 3 – 100 PFU, Group 5 – 1000 PFU and Group 7 – 10,000 PFU). The composition of the vaccine used for the exposure study is described in Supplementary Material Section 3. A second set of three control groups of mice (groups 2, 4 and 6), ten mice per group (mock groups), received PBS buffer solution alone and served as control animals for the study and were similarly challenged with maEBOV. Group 1 animals served as study controls and received no PBS buffer, vaccine or maEBOV injections. All mice were sourced from Jackson Labs and were 6-8 weeks of age and 15-25 grams at the time of vaccination. The dosing regimen is outlined in Table 2.

**Table 2:**
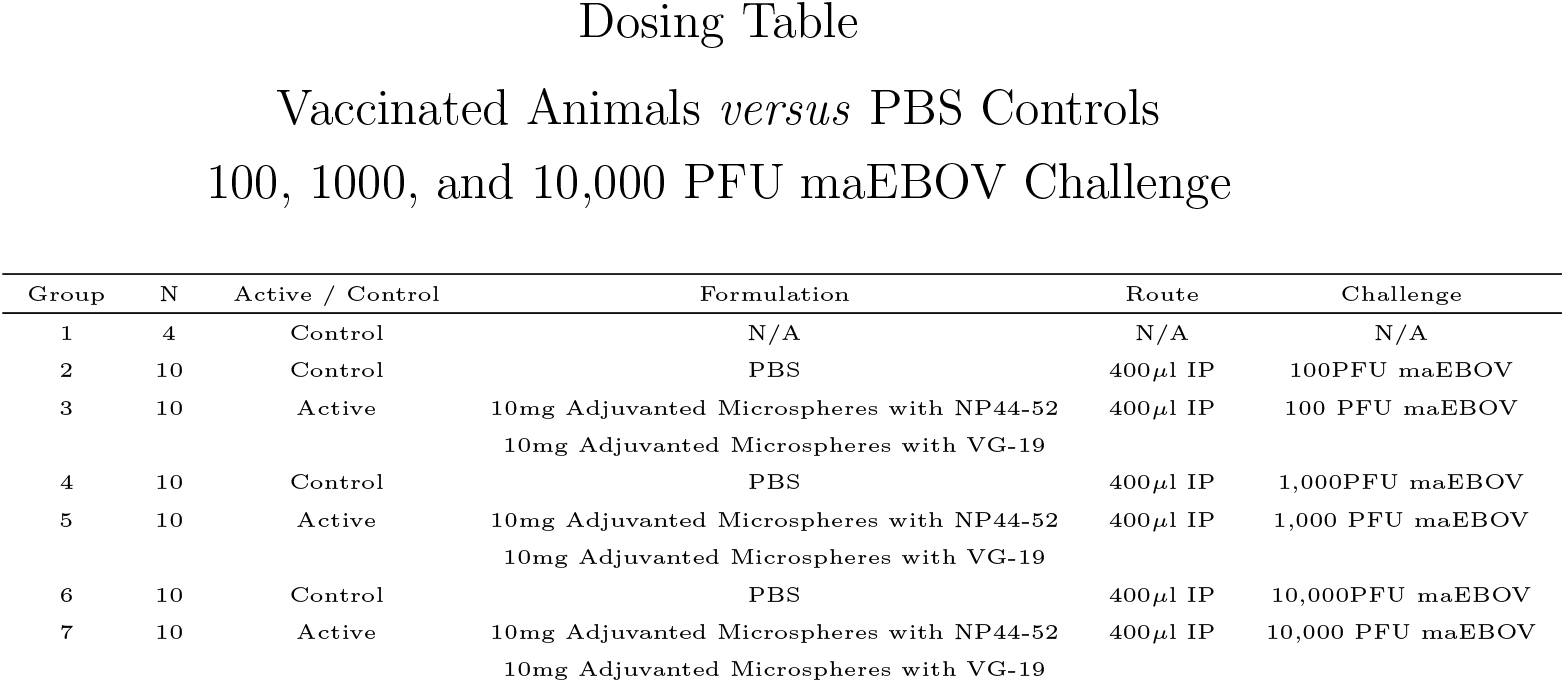
C7BL/6 maEBOV challenge study dosing regimen with PBS (buffer) controls. All challenges were done with Ebola virus M. musculus/COD/1976/Mayinga-CDC-808012 (maEBOV) delivered IP. Mice in Group 1 received no injections.

| Group | N | Active / Control | Formulation | Route | Challenge |
| --- | --- | --- | --- | --- | --- |
| 1 | 4 | Control | N/A | N/A | N/A |
| 2 | 10 | Control | PBS | 400μl IP | 100PFU maEBOV |
| 3 | 10 | Active | 10mg Adjuvanted Microspheres with NP44-52<br>10mg Adjuvanted Microspheres with VG-19 | 400μl IP | 100 PFU maEBOV |
| 4 | 10 | Control | PBS | 400μl IP | 1,000PFU maEBOV |
| 5 | 10 | Active | 10mg Adjuvanted Microspheres with NP44-52<br>10mg Adjuvanted Microspheres with VG-19 | 400μl IP | 1,000 PFU maEBOV |
| 6 | 10 | Control | PBS | 400μl IP | 10,000PFU maEBOV |
| 7 | 10 | Active | 10mg Adjuvanted Microspheres with NP44-52<br>10mg Adjuvanted Microspheres with VG-19 | 400μl IP | 10,000 PFU maEBOV |

Peak mortality across all groups tested was seen in mice challenged with 1,000 PFU maEBOV *versus* PBS buffer control as shown in the survival curve in Figure 3a. Clinical observation data shown in Figure 3b and Figure 3c and daily weight data shown in Figure 3d and Figure 3e show protection from morbidity in all active vaccinated mice exposed to 1,000 PFU maEBOV.

**Figure 3:**
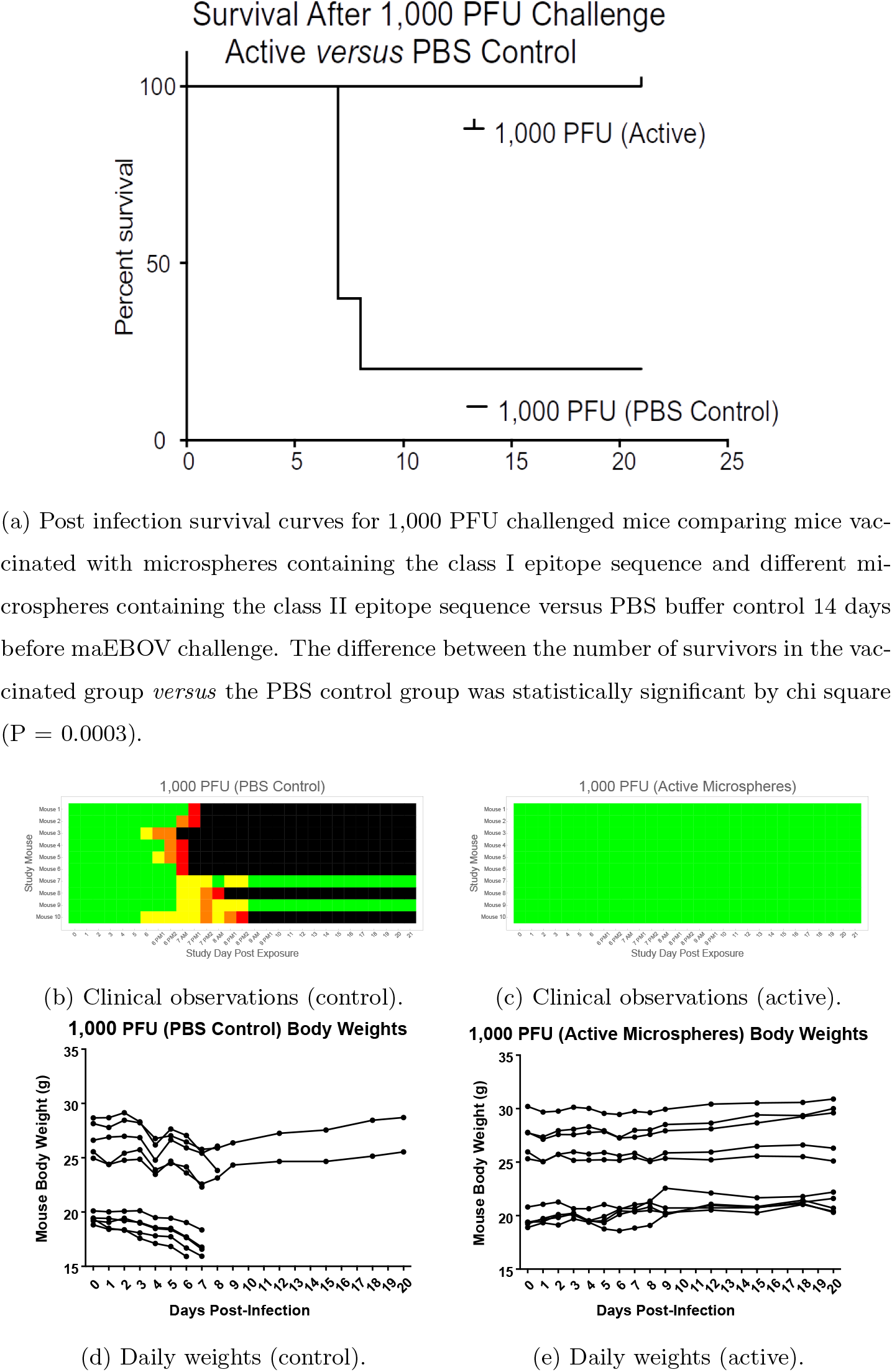
1000 PFU post-challenge data (20mg active adjuvanted microspheres via intraperitoneal injection *versus* PBS buffer solution) collected beginning 14 days after vaccination.

PBS buffer mock-vaccinated mice showed mortality increasing from the 100 PFU to 1,000 PFU as shown in Figure 4a and Figure 3a. We saw a paradoxical effect in control animals with survival increasing between 1,000PFU (Figure 3a) and 10,000 PFU (Figure 5a). We believe this was caused by innate immunity triggered by the very large maEBOV challenge. All mice in all vaccinated groups across both experiments survived and showed no morbidity by clinical observation scores and weight data.

**Figure 4:**
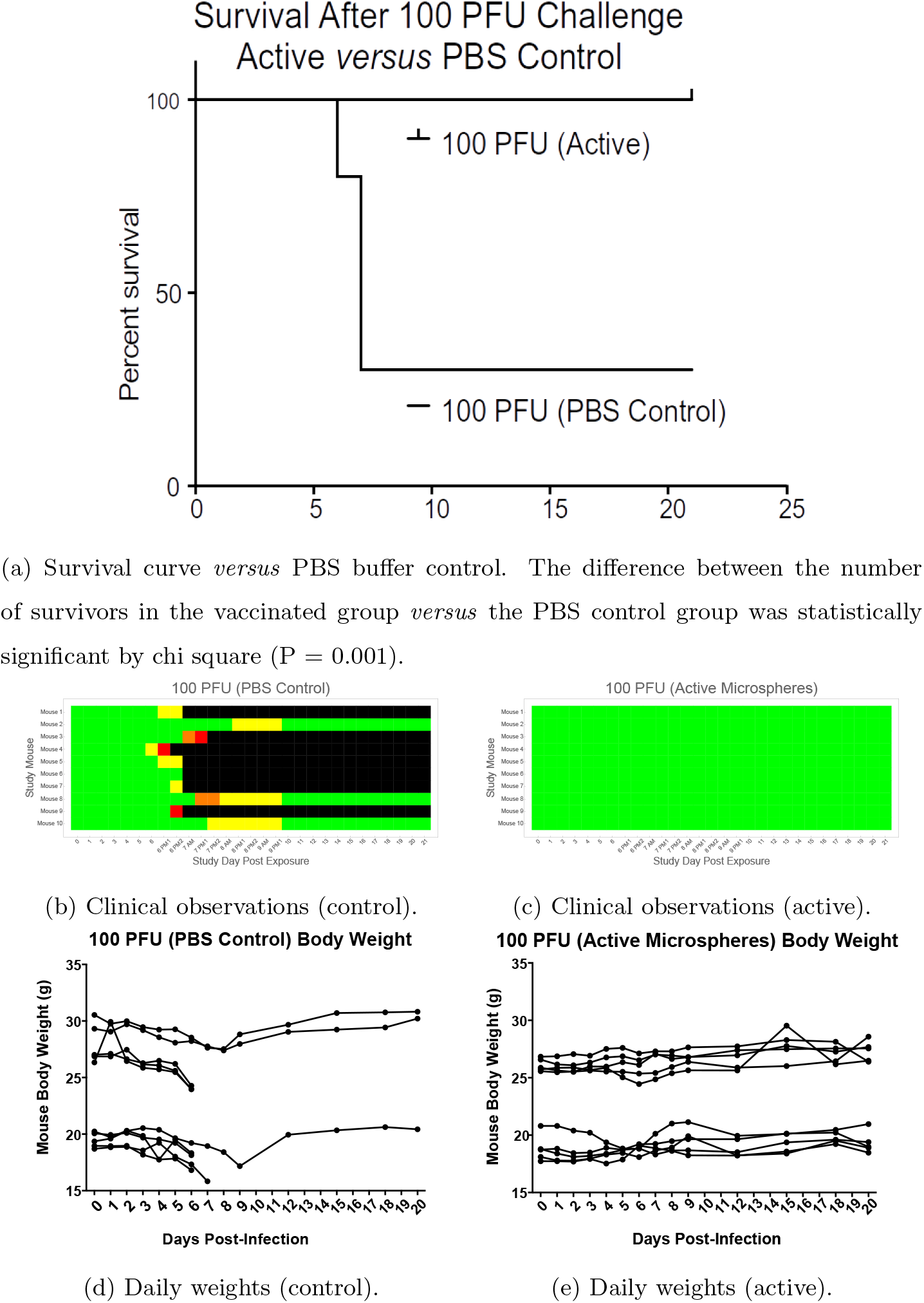
100 PFU post-challenge data (20mg active adjuvanted microspheres via intraperitoneal injection *versus* PBS buffer solution) collected beginning 14 days after vaccination.

**Figure 5:**
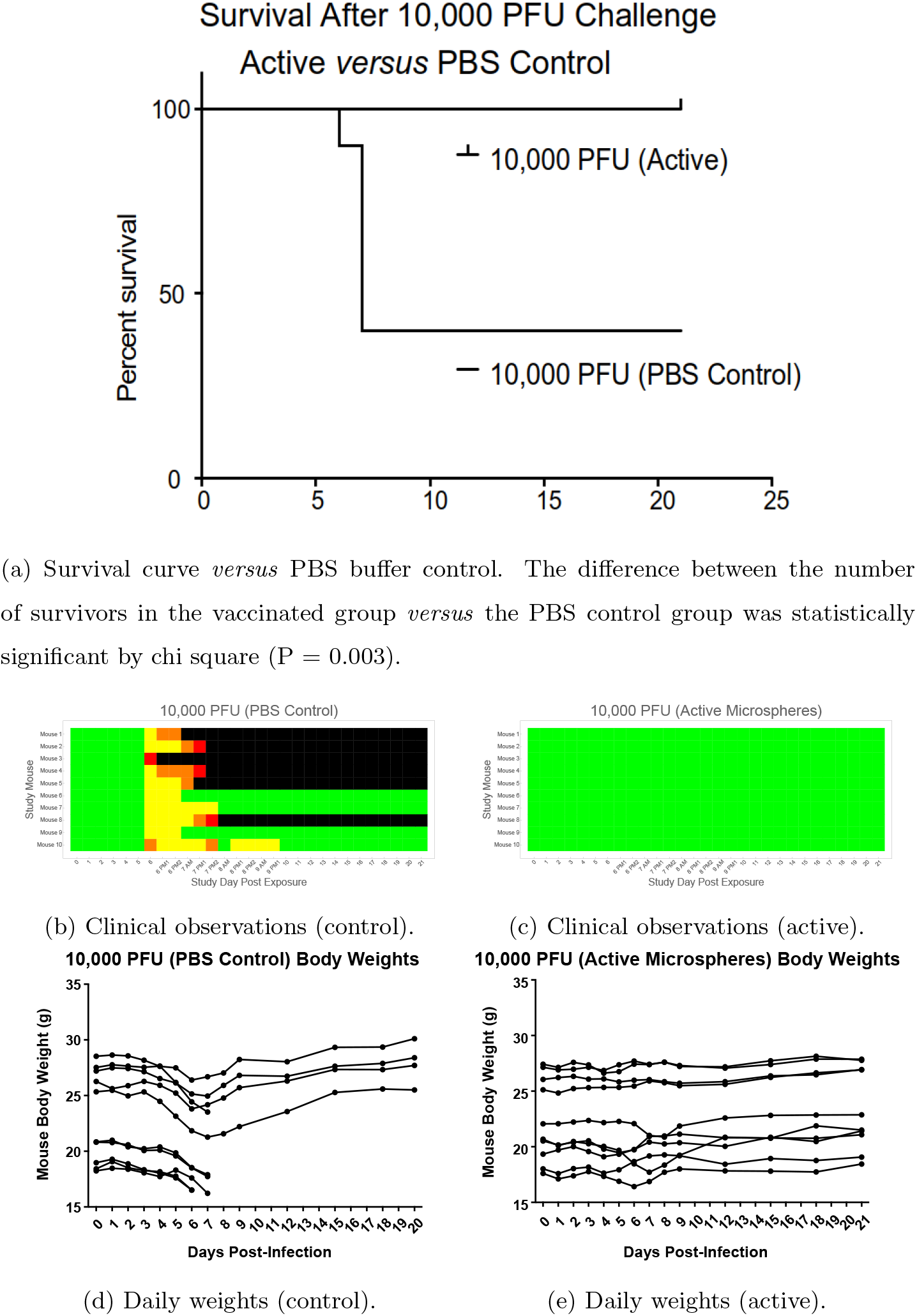
10,000 PFU post-challenge data (20mg active adjuvanted microspheres via intraperitoneal injection *versus* PBS buffer solution) collected beginning 14 days after vaccination.

For each of the three challenge levels, the difference between the number of survivors in the vaccinated group *versus* the PBS control group was statistically significant by chi square (100 PFU P = 0.001; 1000 PFU P = 0.0003; 10,000 PFU P = 0.003).

We saw what appears to be an innate immune response at the 10,000 PFU EBOV exposure level. It has been suggested that EBOV can mediate an innate immunity response through stimulation of TLR-4 [33]. Because the adjuvanted microsphere vaccine used in this experiment incorporates a TLR-4 agonist, we dosed 10 mice with adjuvanted microspheres without peptides and found the level of protection after exposure to 100 PFU EBOV to be statistically no different from that seen in PBS buffer controls (Supplementary Material Figure 1). We conclude that level of protection conferred by the adjuvanted vaccine described in this study is dependant on delivering peptides with the microspheres.

The data in Supplementary Material Figure 1 also shows, in two separate experiments conducted months apart with the same 100 PFU maEBOV challenge dose and the same (active) vaccine formulation, that the vaccinated animals in both active groups had 100% survival and no morbidity by clinical observation. This provides some evidence that the protective effect of vaccination using this adjuvanted microsphere vaccine is reproducible.

Serum samples from sacrificed animals exposed to EBOV who did not receive vaccine were quantitatively assayed for various cytokines using BioPlex plates. Animals having unwitnessed demise did not have serum samples collected. A Pearson Correlation Analysis was performed to assess relationships between specific cytokine levels and survival. The results are shown in Table 3.

**Table 3:**
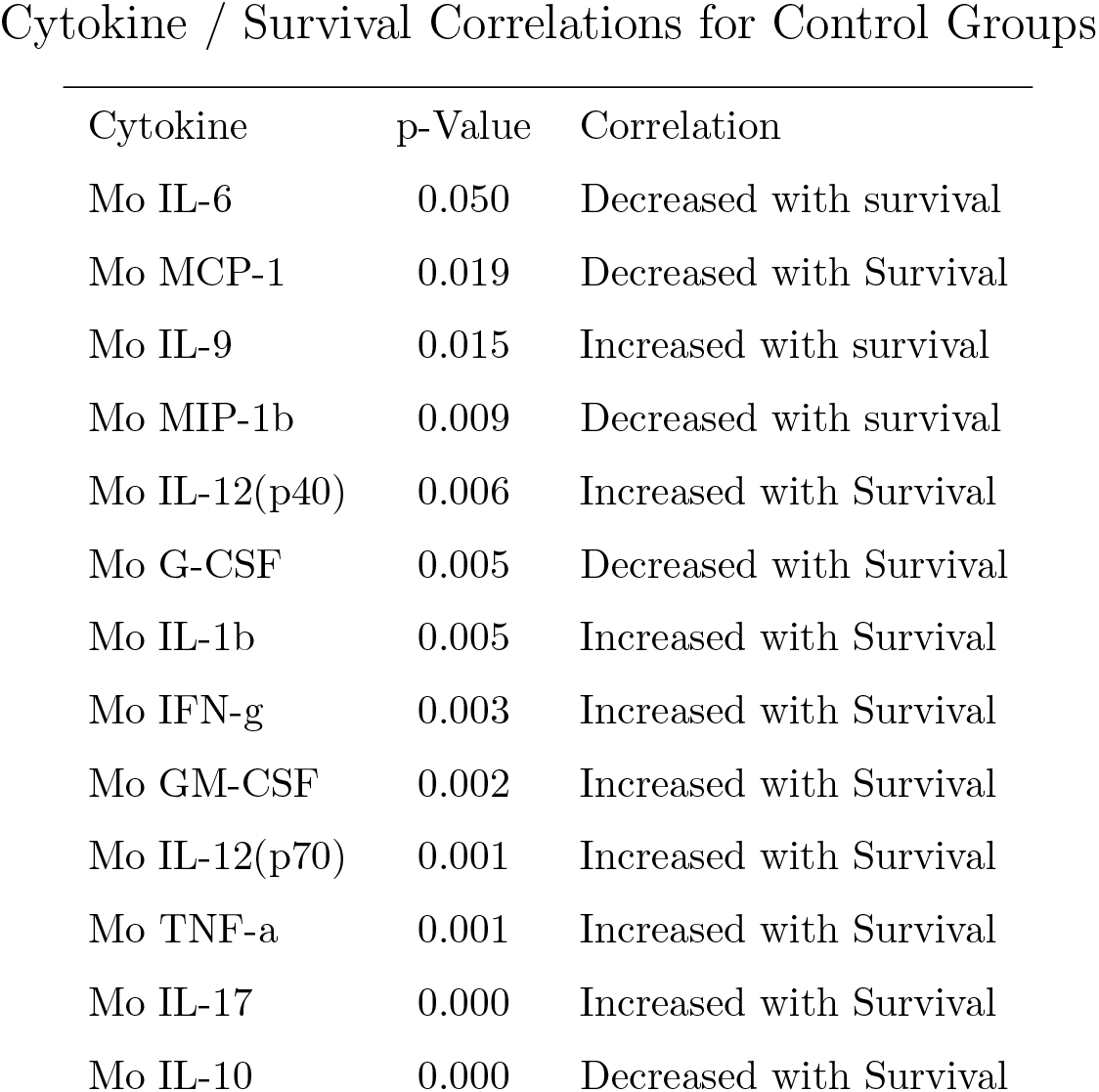
Cytokines with statistically significant (positive or negative) correlation with survival in non-vaccinated mice are shown here along with (Pearson Correlation Analysis) p-values.

We observed low levels of IL-6 in surviving mice. NHPs infected with EBOV have been determined by other researchers to have elevated levels of IL-6 in plasma and serum [27][17]. EBOV infected humans have also shown elevated IL-6 levels and these elevated levels have been associated with increased mortality [71].

Similarly, we observed low levels of MCP-1, IL-9 and GM-CSF in survivors. Increased serum and plasma levels of MCP-1 have been observed in EBOV infected NHPs [22][27][17] and elevated levels of MCP-1 were associated with fatalities in EBOV infected human subjects [71]. Human survivors of EBOV have been found to have very low levels of circulating cytokines IL-9 and elevated levels of GM-CSF have been associated with fatality in humans exposed to EBOV [71].

We saw increased levels of *IFN — γ* in survivors. Other vaccine studies have associated *IFN —* γ with protection [70][38].

We achieved protection against maEBOV challenge with a single injection of an adjuvanted microsphere peptide vaccine loaded with a class I peptide in a region on EBOV nucleocapsid favored for CD8+ attack by survivors of the 2013-2016 West Africa EBOV outbreak. There is evidence that a CTL response could be beneficial in the context of a coronavirus infection. [13][41][64][11][28][36] Peng et al. have found survivors of the SARS-CoV-1 outbreak who had circulating T-cells targeting SARS-CoV-1 nucleocapsid two years after initial infection. [42] We decided to investigate the feasibility of designing a SARS-CoV-2 peptide vaccine targeting SARS-CoV-2 nucleocapsid.

All available SARS-CoV-2 protein sequences were obtained from the NCBI viral genomes resource within GenBank, an NIH genetic sequence database [8]. Retrieved sequences were processed using multiple sequence alignment (MSA) via Clustal for the nucleocapsid phosphoprotein [34]. The nucleocapsid phosphoprotein sequences were trimmed down to every possible peptide sequence 9 amino acids in length. 9mers were chosen because they typically represent the optimal length for binding to the vast majority of HLA [1][18][3]. The resulting peptides were compared to the MSA to ensure than these sequences are conserved within all of the sequencing samples available and not affected by an amino acid variant that could complicate subsequent analysis, specifically the calculation of population coverage. A selection of HLA were selected to encompass the vast majority of the worlds population at over 97% coverage.

Peptides were run through artificial intelligence algorithms, netMHC and netMHCpan which were developed using training data from *in-vitro* binding studies. The pan variant of netMHC is able to integrate *in-vitro* data from a variety of HLA to allow for predictions to be made if limited *in-vitro* data is available for the specified target HLA [30][1]. This *in-silico* analysis utilizes the neural networks ability to learn from the *in-vitro* data and report back predicted values based on the imputed SARS-CoV-2 nucleocapsid phosphoprotein peptides. Peptides with a predicted HLA IC50 binding affinity of 500nm or less in either of the algorithms, were included in the candidate list of targets for the vaccine [30][1][40].

A subset of these SARS-CoV-2 peptide sequences are present on SARS-CoV-1 nucleocapsid phosphoprotein and as a result had in-vitro binding data in Immunology Epitope Database and Analysis Resource (IEDB) collected after a previous outbreak [65]. Predicted values of these peptides were cross referenced with actual *in-vitro* binding measurements from identical 9mer peptides when that data was available.

## 3. Summary and Discussion

Most preventative vaccines are designed to elicit a humoral immune response, typically via the administration of whole protein from a pathogen. Antibody vaccines typically do not produce a robust T-cell response. [72] A T-cell vaccine is meant to elicit a cellular immune response directing CD8+ cells to expand and attack cells presenting the HLA Class I restricted pathogen-derived peptide antigen. [47] Difficulty in obtaining a reliable immune response from peptide antigens and the HLA restricted nature of CTL vaccines have limited their utility to protect individuals from infectious disease [77]. However, observations derived from individuals able to control HIV infection [43] and EBOV infection [50] demonstrating that control may be associated with specific CTL targeting behavior, suggest that there may be an important role for HLA-restricted peptide vaccines for protection against infectious disease for which development of an effective traditional whole protein vaccine has proved to be difficult. The adjuvanted microsphere peptide vaccine platform described here incorporates unmodified peptides making possible rapid manufacture and deployment to respond to a new viral threat.

NP44-52 is located within one of the EBOV nucleocapsid proteins considered essential for virus replication. This epitope resides in a sequence conserved across multiple EBOV strains as shown in Supplementary Material Figure 6. A 7.3Å structure for NP and VP24 is shown for context in Figure 6a [67]. A 1.8Å resolution structure rendering for EBOV NP shown in Figure 6b illustrates that NP44-52 is a buried structural loop, which is likely to be important to the structural integrity of the EBOV NP protein [16]. This structural role of NP44-52 likely explains its conservation across EBOV strains.

**Figure 6:**
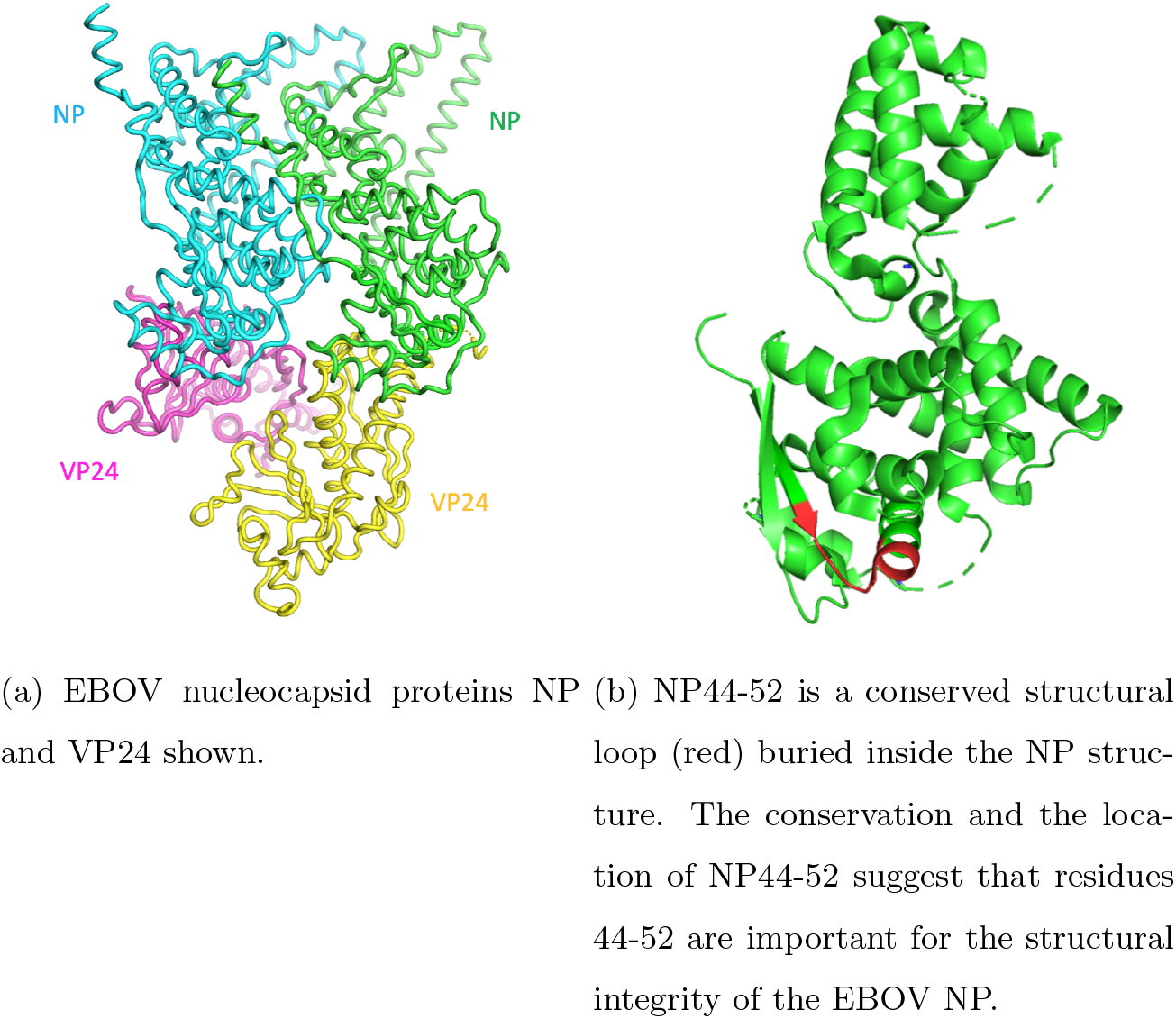
The class I epitope used for this study is located within NP. Nucleocapsid proteins NP and VP24 are shown together in (a). A detailed view of NP with the study epitope position highlighted in shown in (b).

CTL targeting of the EBOV NP protein has been described [42][64][28] [49][24]. Nucleocapisid proteins are essential for EBOV replication [61]. Recent advances in T-cell based vaccines have focused on avoiding all variable viral epitopes and incorporating only conserved regions [7][25]. EBOV NP may be more conserved than nucleocapsid proteins VP35 and VP24 making it more suitable as a CTL vaccine target [9][73]. The nucleocapsid proteins in SARS-CoV-1 are also essential for that virus to function normally [10]. This suggests that a CTL vaccine targeting coronavirus nucleocapsid could be effective against SARS-CoV-1 or SARS-CoV-2.

We have shown that an H2-D^b^ restricted Class I peptide exists within the NP41-60 epitope identified by Sakebe et al. as the most commonly favored NP epitope for CD8+ attack by survivors of the 2013-1016 EBOV outbreak in West Africa. We have demonstrated, when delivered in conjunction with a predicted-matched Class II epitope using an adjuvanted microsphere peptide vaccine platform, NP44-52 protection against mortality and morbidity for the maEBOV challenge doses tested in a C57BL/6 mouse model. We accomplished this with an adjuvanted, microsphere-based, synthetic CTL peptide vaccine platform producing a protective immune response 14 days after a single administration.

EBOV can cause severe pulmonary problems in exposed subjects [39]. These problems can be especially severe when the virus is delivered by aerosol [15][31]. Interaction of EBOV specific antibody, NHP lung tissue and EBOV delivered to NHPs via aerosol can produce a more lethal effect than in NHPs without circulating anti-EBOV antibody exposed to aerosolized EBOV (unpublished conference presentation). This suggests that a CTL vaccine may be more effective for prophylaxis against filovirus protection than an antibody vaccine if the anticipated route of EBOV exposure is via aerosol.

Sakebe et al. identified A*30:01:01 as the only HLA type common to all four survivors in their study with CD8+ targeting of NP41-60. The A*30 supertype is relatively common in West Africa: 13.3% for Mali, 15.4% for Kenya, 16.3% for Uganda, and 23.9% for Mozambique [32]. Although peptide vaccines are by their nature HLA restricted, it may be possible to create a CTL vaccine directed against EBOV for use alone or in conjunction with a whole protein vaccine to produce an antibody response in tandem, by incorporating additional Class I peptides from epitopes targeted by controllers to broaden the HLA coverage of the vaccine. MHC binding algorithms hosted by the IEDB predict that YQVNNLEEI will bind strongly to the MHC of HLA-A*02:06, HLA-A*02:03 and HLA-A*02:01 individuals (Supplementary Material Table 2) [65]. HLA-DR binding database analysis also suggests that VKNEVNSFKAALSSLAKHG demonstrates sufficiently promiscuous binding characteristics cover that same population (Supplementary Material Table 4) [65]. Taken together, a peptide vaccine based on YQVNNLEEI and VKNEVNSFKAALSSLAKHG could produce a cellular immune response in about 50% of the population of the Sudan and about 30% of the population of North America.

The internal proteins located within influenza virus, in contrast to the glycoproteins present on the surface, show a high degree of conservation. Epitopes within these internal proteins often stimulate T-cell-mediated immune responses [57]. As a result, vaccines stimulating influenza specific T-cell immunity have been considered as candidates for a universal influenza vaccine [66].

SARS-CoV-1 infection survivors have been found to have a persistent CTL response to SARS-CoV-1 nucleocapsid two years after infection. [42] This suggests that the same approach could be applied to SARS-CoV-2 which has conserved regions in nucleocapsid which is located within the virus (see multiple sequence alignment in Supplementary Material Figure 7 and Supplementary Material Figure 8). Antigenic escape allows a virus to retain fitness despite an immune response to vaccination [20]. Picking conserved regions for vaccine targeting is an important part of mitigating this problem. Coronavirus spike protein, for example, may be particularly susceptible to mutation meaning that antigenic escape would be likely if the spike protein was targeted by a coronavirus vaccine, making it difficult to achieve durable protection. [74] A recent paper conducted a population genetic analysis of 103 SARS-CoV-2 genomes showing that the virus has evolved into two major types: L and S, with changes in their relative frequency after the outbreak possibly due to human intervention resulting in selection pressure [62].

We took all possible 424 9mer peptide sequences from the SARS-CoV-2 nucleocapsid protein sequences available and evaluated each peptide for HLA restriction using NetMHC 4.0 and NetMHCpan 4.0 [65][30][1]. We analyzed 9mer peptide sequences because these are often associated with superior MHC binding properties than class I peptides of other lengths [63][18]. We found 53 unique peptides with predicted binding below 500nM from NetMHC 4.0 and/or NetMHCpan 4.0. These results are shown in Supplementary Material Table 6, Supplementary Material Table 7, Supplementary Material Table 8 and Supplementary Material Table 9.

We proceeded to determine the predicted HLA population coverage of a vaccine incorporating all 53 peptides using median values of the ANN, SMM, NetMHC 4.0 and NetMHCpan 4.0 algorithms hosted by IEDB [65]. These 53 peptides, taken together, had predicted HLA coverage of greater than 97% of the world’s population as shown in Supplementary Material Table 10. We also calculated HLA coverage based on alleles specific to populations in China and found that coverage across those individuals could be expected to be within 3% percent of the world wide coverage estimate as shown in Supplementary Material Table 11. This same population coverage could be achieved with 16 of the 53 unique peptides as shown in Table 4.

**Table 4:**
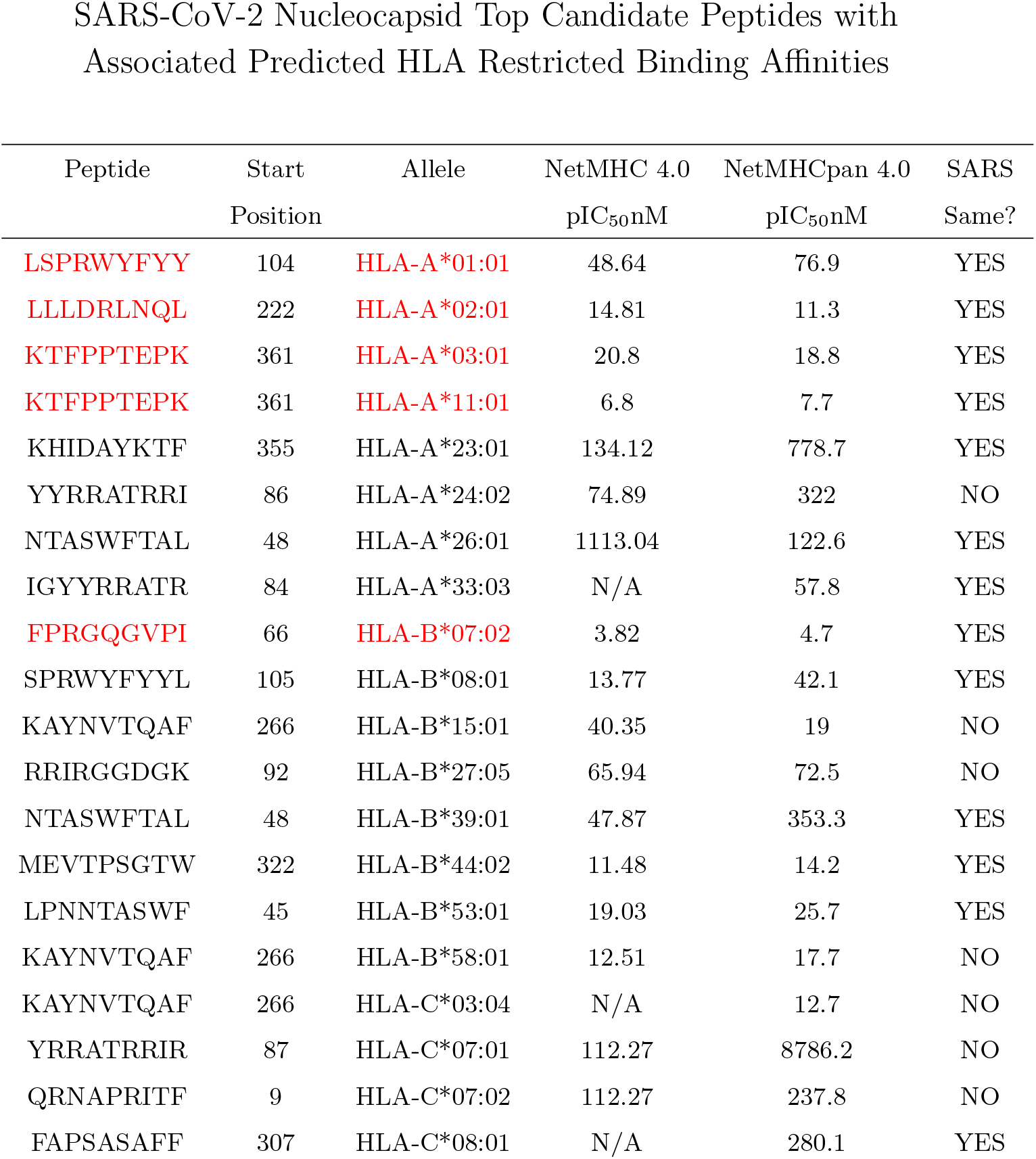
This set of 16 unique peptides represents the minimum number required to achieve > 95% world-wide population coverage. The starting position is within the nucleocapsid. Top binding affinity predictions chosen via NetMHC 4.0 or NetMHCpan 4.0. Peptide sequences colored in red have literature references as known *in-vitro* binders to the predicted allele match (see text).

Seven of the 53 peptides with a predicted HLA match have been tested *in-vitro* for HLA binding affinity by various researchers [65]. These binding affinity assays were originally performed with the SARS virus during a previous outbreak. Specific literature references for these *in-vitro* assays for each peptide sequence are as follows: ASAFFGMSR, LSPRWYFYY, QQQGQTVTK: [53], FPRGQGVPI: [53][26][46][60], GMSRIGMEV: [26][64][13][41][12], KTF-PPTEPK: [53][26][45][60][6] and LLLDRLNQL: [41][13][12][64][78]. These seven peptides are shown in red in Supplementary Material Table 6 and Supplementary Material Table 7.

The remaining 46 SARS-CoV-2 peptides listed in could also be further qualified as potential vaccine candidates by confirming MHC binding predictions by *in-vitro* binding affinity and/or binding stability studies [54][52][26]. Another approach to evaluating the 53 SARS-CoV-2 candidate vaccine peptides though *in-vitro* testing is also possible.

As we have shown in this paper, a peptide targeted by EBOV controllers could form the basis of a preventative vaccine for EBOV. ELISPOT analysis of PBMCs taken from the peripheral blood of COVID-19 controllers and progressors to assess the presence of a differential response to the 53 peptides could lead to a broadly applicable protective CTL vaccine against SARS-CoV-2 by incorporating peptides into the vaccine that are more commonly targeted for CD8+ attack by the controllers *versus* the progressors. A peptide vaccine for SARS-CoV-2, unlike a typical antibody vaccine, is not limited to virus surface antigen targets. This provides opportunities to attack other targets on SARS-CoV-2 besides spike which may be prone to mutation [74].

In addition, a peptide vaccine mitigates the risk of Antibody Disease Enhancement (ADE) seen in the context of a non-neutralizing antibody response to a whole protein vaccine [75][55]. Also, neutralizing antibodies directed against spike protein in SARS-CoV-1 patients have been associated with an increased risk of Acute Lung Injury (ALI)[35]. Specifically, patients succumbing to SARS-CoV-1 were found to develop a neutralizing antibody (NAb) response to spike protein faster than survivors after the onset of symptoms and the NAb titers were higher in the patients who died compared with those who recovered[76]. To the extent to which antibody vaccines producing an antibody response against the spike protein in SARS-CoV-2 could increase the risk of ALI, this risk could also be mitigated by a using peptide vaccine as an alternative approach.

The extent of the COVID-19 outbreak should allow many more controllers to be identified than the thirty individuals studied by Sakabe and the seven individuals identified in the Peng study [42][50]. Furthermore, Sakebe and Peng did not report progressor data perhaps because of the difficulty in obtaining blood samples from those patients. If researchers act now during the COVID-19 outbreak, perhaps controller and progressor blood samples could be collected and prospectively analyzed, quickly creating a database of optimal candidate class I peptides for inclusion into a CTL vaccine with potentially broad HLA coverage for subsequent rapid manufacture and deployment. It would be interesting to see the extent to which the peptides favored by controllers appear on SARS-CoV-2 nucleocapsid, making SARS-CoV-2 a second example, across two different viruses, of controllers exhibiting CTL attack preferentially on the nucleocapsid protein.

## Supporting information

ebola_supplementary

## 4. Acknowlegements

All animal handling was done in accordance with NIH and institutional animal care and use guidelines by Aragen Bio-sciences in Morgan Hill California and the University of Texas, Medical Branch, Galveston Texas working in conjunction with the Galveston National Laboratory. The research was funded by Flow Pharma, Inc.

