## Supplementary material for "An Effective CTL Peptide Vaccine for Ebola Zaire Based on Survivors’ CD8+ Targeting of a Particular Nucleocapsid Protein Epitope with Potential Implications for COVID-19 Vaccine Design": ebola_supplementary

### 1. Mitigating Potential for Epitope Competitive Inhibition at the MHC

The possibility of competitive inhibition at the MHC when a CTL vaccine containing multiple peptides is administered to a patient has been acknowledged [4]. Techniques such as splitting a CTL vaccine dose across multiple injection sites with one peptide sequence per site have been suggested to address this issue [8]. We demonstrated *in-vitro* that competitive inhibition at the MHC could occur by incubating 200,000 PBMCs per well from a reference PMBC sample (CTL Reference Sample QC Set, Cellular Technology, Ltd. Shaker Heights, Ohio), from an HLA A*02, HIV naive subject with an HCMV Class I epitope pp65 (495-503) NLVPMVATV supplied with the Reference Sample and known to produce $IFN-\gamma$ release in that reference PBMC sample and then introducing increasing concentrations of of an HLA A*02 matched class I HIV peptide. Figure 9 shows functional inhibition of MHC binding as evidenced by decreasing, and ultimate extinction of $IFN-\gamma$ release as determined by ELISA assay with increasing concentrations of the HLA matched HIV peptide. This experiment suggests that competitive inhibition could occur when two different HLA matched peptide sequences are delivered to the same antigen presenting cell. The adjuvanted microspheres used in this study were manufactured to be nominally the same same size as antigen presenting cells (about 11$\mu$M). By loading each peptide microsphere with only one peptide sequence, this mitigates against two peptides being processed by the same antigen presenting cell at the same time. Multiple peptides can be incorporated into a vaccine formulation by blending different populations of adjuvanted microspheres into the same vaccine dose, with each microsphere containing only one peptide sequence.

### 2. Route of Administration

Intradermal injection of influenza vaccine has been shown to be more effective than intramuscular administration in human subjects [3]. Gamma scintigraphy studies have shown that the efficacy of intradermally-delivered antigen may be due to the portion of antigen that reaches the lymph system [9]. The rat peritoneal space has been demonstrated to drain into the rat lymphatics [5]. We sought to determine if H2D-Kb matched Class I epitopes delivered into the mouse intraperitoneal space with Class II eptiopes would provide a stronger immune response by ELISPOT-determined $IFN-\gamma$ release compared with intradermal and intramuscular administration to C57BL/6 mice.

Prior to conducting our challenge study, we used adjuvanted microspherers to immunize C57BL/6 mice [6]. OVA and VSV class I and class II epitopes known to produce an immune response in this mouse model were selected and delivered by three different routes of administration [2] [10] [7] [1]. Four different microsphere populations, each containing CpG and one of the epitopes from Table 14, were prepared. The four microsphere populations were blended 1:1:1:1 by weight and suspended just prior to administration in a PBS injectate solution containing MPLA. A total of 2mg of the microsphere preparation was administered into C57BL/6 mice by the intradermal, base of tail (ID), intramuscular (IM) or intraperitoneal (IP) route using four mice for each route of administration. Splenocytes were harvested on day 14 and subjected to ELISPOT analysis evaluating $IFN-\gamma$ release in response to stimulation with the peptide used in the vaccination. As illustrated in Figure 11 and Figure 10, the CTL response to each of the administered class I epitopes was significantly higher for the IP route of administration. Guided by this data, we selected IP as the route of vaccine administration for our EBOV challenge study.

#### 3. Vaccine Formulation Composition

Adjuvanted microspheres nominally 11$\mu$M in diameter (geometric standard deviation 1.2) manufactured as a room temperature stable dry powder and loaded as described in Table 12 are mixed with the injectate solution described in Table 13 just prior to injection. All mice in the EBOV challenge study received nominally 450$\mu$l of this adjuvanted microsphere suspension by intraperitoneal administration.

#### 4. Separation of CD4 & CD8 T-cells for ELISPOT

Mouse spleens were harvested and splenocytes were obtained by standard procedures. Splenocytes were counted and divided into three equal groups; one for total cell response in the ELISPOT assay, one for the ELISPOT response in the absence of CD4 cells, and the final group for the ELISPOT response in the absence of CD8 cells. CD4 and CD8 cells were removed by positive selection using magnetic beads coated with either anti-CD4 or anti-CD8 antibody respectively, and a MACS Separator (Miltenyi Biotech, Auburn, CA) according to the manufacturer's instructions. Cells not bound to the columns (i.e., unlabeled by the specific antibody) were collected, centrifuged, washed, re-centrifuged, counted, and placed in a standard ELISPOT assay as previously described.

Mouse Observation Clinical Scores

| Scale | Description of Animal |
| --- | --- |
| 1 | Healthy |
| 2 | Lethargic and/or ruffled fur |
| | (triggers a second observation) |
| 3 | Ruffled fur, lethargic and hunched posture, orbital tightening |
| | (triggers a third observation) |
| 4 | Ruffled fur, lethargic, hunched posture, orbital tightening |
| | reluctance to move when stimulated, paralysis or greater than 20% weight loss |
| | (requires immediate euthanasia) |

Table 1: Clinical score indices used to track morbidity in study animals.

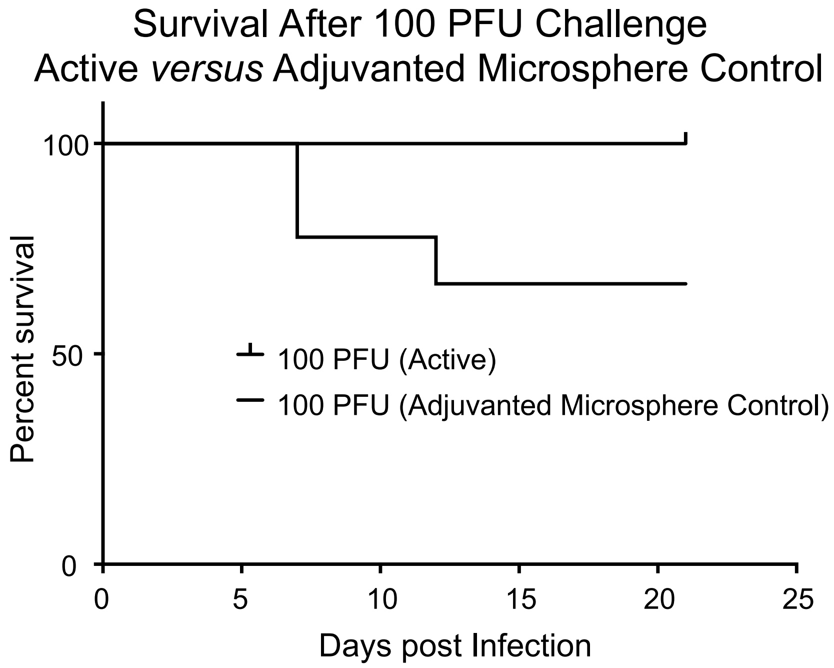

(a) Survival curve *versus* Adjuvanted Microsphere Control.

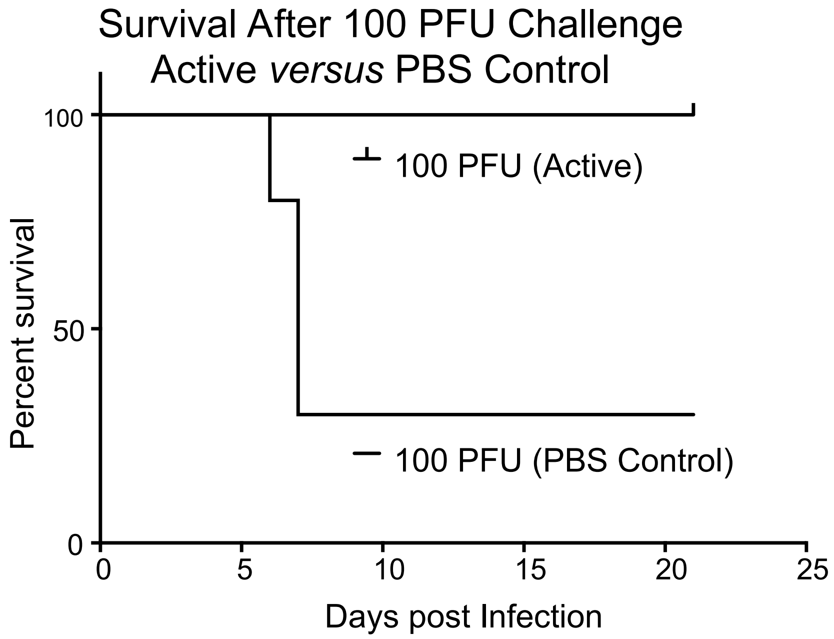

(b) Survival curve *versus* PBS buffer control.

Figure 1: The 100PFU PBS control survival curve 1b is not statistically different from the 100PFU adjuvanted control survival curve 1a. A chi square test results in a P = 0.37 for (survived / dead) 6/4 (adjuvanted) versus 3/7 (PBS control).

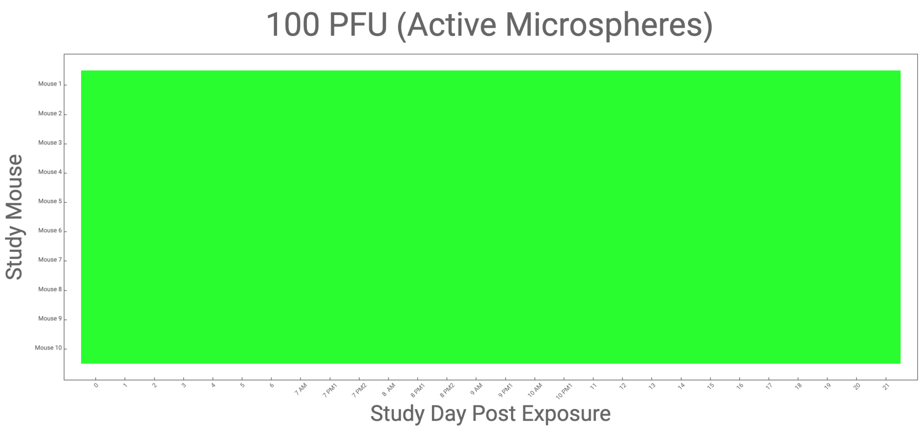

Figure 2: Clinical observations showing that no mice died in this group during the study, scored from 1 (healthy) to 4 (moribund) made post infection in control animals receiving PBS buffer via intraperitoneal injection 14 days before infection. The clinical scores described in Table 1 are shown using the following color scheme: 1 = GREEN, 2 = YELLOW, 3 = ORANGE and 4 = RED. A dead mouse is coded in black. The frequency of measurements was increased on post infection days 6-9 coinciding with the anticipated period of peak morbidity.

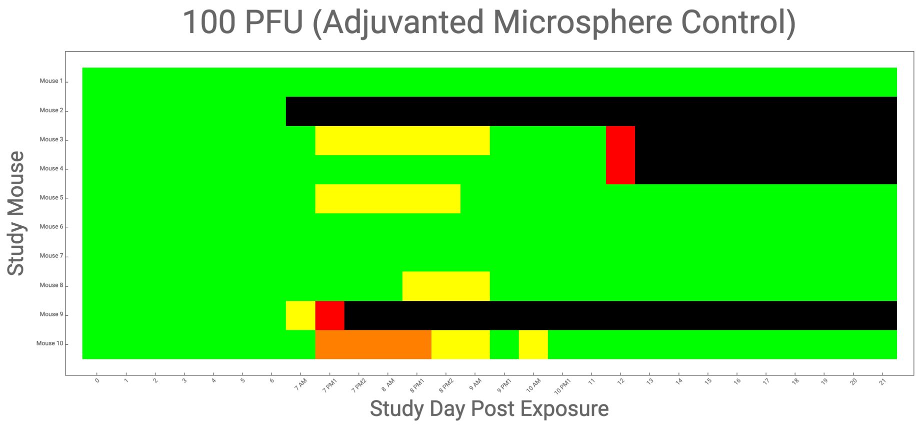

Figure 3: Clinical observations, scored from 1 (healthy) to 4 (moribund) made post infection in control animals receiving PBS buffer via intraperitoneal injection 14 days before infection. The clinical scores described in Table 1 are shown using the following color scheme: 1 = GREEN, 2 = YELLOW, 3 = ORANGE and 4 = RED. A dead mouse is coded in black. The frequency of measurements was increased on post infection days 6-9 coinciding with the anticipated period of peak morbidity.

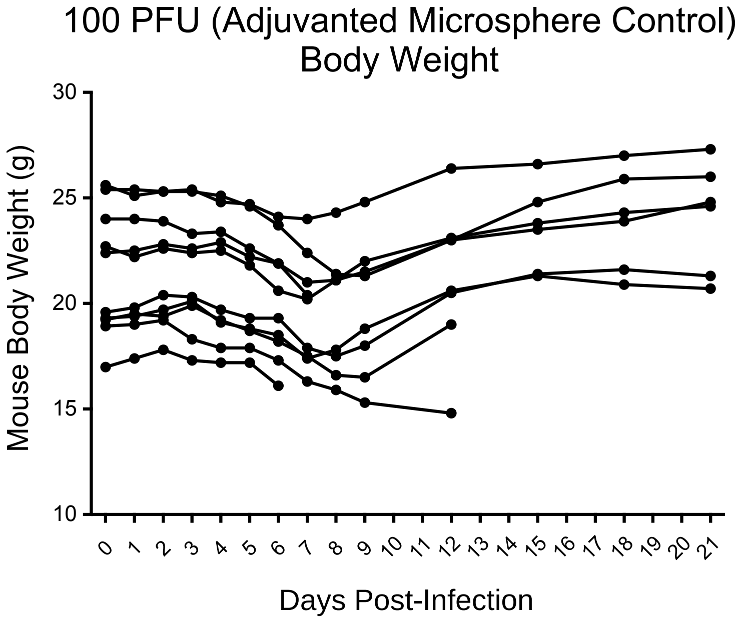

Figure 4: Daily weights were recorded post infection. Measurements for control animals, receiving adjuvanted microspheres 14 days before infection, are shown here.

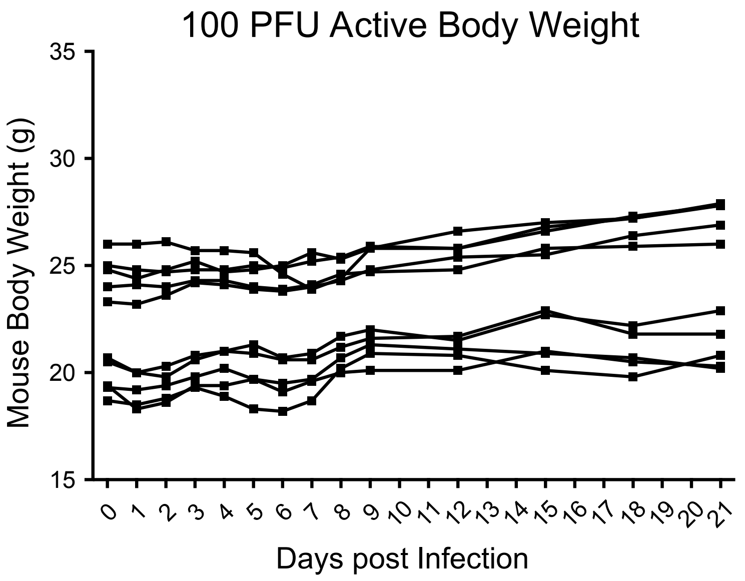

Figure 5: Daily weights were recorded post infection. Measurements for animals receiving active vaccine 14 days before infection, are shown here.

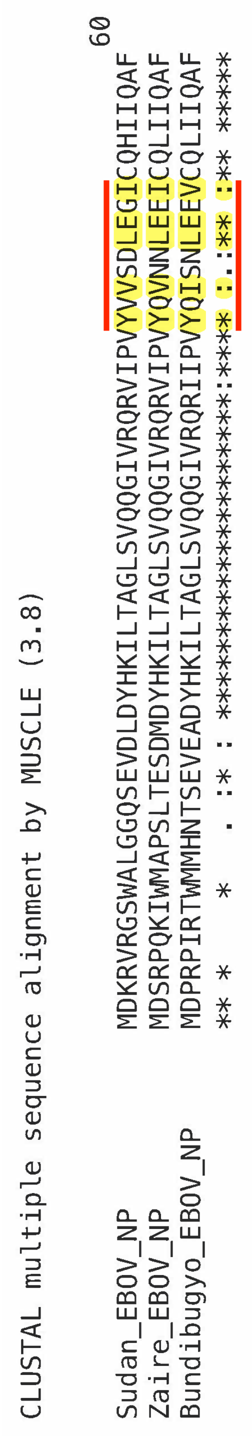

Figure 6: NP44-52 has conserved residues across three strains of EBOV.

Computed HLA Binding Affinities for YQVNNLEEI

| Allele | Median $pIC_{50}nM$ | Consensus Score |
|---|---|---|
| **HLA-A*02:06** | 5.8 | 0.16 |
| **HLA-A*02:03** | 106 | 2.4 |
| **HLA-A*02:01** | 198 | 3.7 |
| HLA-B*15:01 | 791 | 4.7 |
| HLA-A*23:01 | 1140 | 2.1 |
| HLA-B*40:01 | 1140 | 2.4 |
| HLA-A*68:02 | 5553 | 23 |
| HLA-A*24:02 | 5664 | 5.7 |
| HLA-B*53:01 | 8737 | 15 |
| HLA-B*58:01 | 12128 | 17 |
| HLA-B*51:01 | 13551 | 6.3 |
| HLA-A*26:01 | 15442 | 11 |
| HLA-A*32:01 | 17173 | 22 |
| HLA-B*44:03 | 18798 | 17 |
| HLA-B*35:01 | 19374 | 35 |
| HLA-A*30:01 | 23549 | 61 |
| HLA-B*44:02 | 27488 | 20 |
| HLA-A*30:02 | 31424 | 55 |

Table 2: Database-predicted HLA binding affinities for NP44-52 (YQVNNLEEI), the class I peptide used in this study.

Computed H-2 Binding Affinities for YQVNNLEEI

| Allele | Median $pIC_{50}nM$ | Consensus Score |
|---|---|---|
| **H-2-D$^{b}$** | 26 | 0.20 |
| H-2-K$^{d}$ | 5639 | 7.0 |
| H-2-K$^{b}$ | 13722 | 32 |
| H-2-D$^{d}$ | 21052 | 23 |
| H-2-L$^{d}$ | - | 41 |

Table 3: Database-predicted H-2 binding affinities for NP44-52 (YQVNNLEEI), the class I peptide used in this study.

### Computed HLA-DR Binding Affinities for VKNEVNSFKAALSSLAKHG

| Allele | Start | 15-mer peptide | Median $pIC_{50}$nM | Consensus Score |
|---|---|---|---|---|
| **HLA-DRB1*01:01** | 5 | VNSFKAALSSLAKHG | 4.1 | 0.28 |
| **HLA-DRB1*09:01** | 5 | VNSFKAALSSLAKHG | 6.0 | 0.010 |
| **HLA-DRB1*04:05** | 4 | EVNSFKAALSSLAKH | 14 | 0.19 |
| **HLA-DRB5*01:01** | 5 | VNSFKAALSSLAKHG | 24 | 1.8 |
| **HLA-DQA1*05:01/DQB1*03:01** | 3 | NEVNSFKAALSSLAK | 24 | 13 |
| **HLA-DRB1*04:01** | 5 | VNSFKAALSSLAKHG | 25 | 1.2 |
| **HLA-DPA1*02:01/DPB1*14:01** | 4 | EVNSFKAALSSLAKH | 27 | 4.7 |
| **HLA-DRB3*02:02** | 4 | EVNSFKAALSSLAKH | 27 | 0.080 |
| **HLA-DRB1*07:01** | 2 | KNEVNSFKAALSSLA | 40 | 4.7 |
| **HLA-DRB1*11:01** | 5 | VNSFKAALSSLAKHG | 66 | 4.3 |
| **HLA-DRB1*15:01** | 3 | NEVNSFKAALSSLAK | 152 | 3.4 |
| **HLA-DRB1*08:02** | 3 | NEVNSFKAALSSLAK | 162 | 1.4 |
| **HLA-DQA1*01:02/DQB1*06:02** | 3 | NEVNSFKAALSSLAK | 167 | 6.2 |
| **HLA-DPA1*02:01/DPB1*01:01** | 4 | EVNSFKAALSSLAKH | 401 | 26 |
| **HLA-DRB1*12:01** | 5 | VNSFKAALSSLAKHG | 769 | 8.8 |
| **HLA-DPA1*03:01/DPB1*04:02** | 4 | EVNSFKAALSSLAKH | 773 | 15 |
| **HLA-DRB4*01:01** | 3 | NEVNSFKAALSSLAK | 903 | 37 |
| HLA-DRB1*13:02 | 1 | VKNEVNSFKAALSSL | 1380 | 28 |
| HLA-DRB1*03:01 | 2 | KNEVNSFKAALSSLA | 1498 | 13 |
| HLA-DQA1*03:01/DQB1*03:02 | 1 | VKNEVNSFKAALSSL | 1680 | 21 |
| HLA-DPA1*02:01/DPB1*05:01 | 5 | VNSFKAALSSLAKHG | 1811 | 25 |
| HLA-DQA1*04:01/DQB1*04:02 | 4 | EVNSFKAALSSLAKH | 1951 | 16 |
| HLA-DRB3*01:01 | 2 | KNEVNSFKAALSSLA | 1991 | 21 |
| HLA-DPA1*01:03/DPB1*02:01 | 4 | EVNSFKAALSSLAKH | 2002 | 26 |
| HLA-DPA1*01/DPB1*04:01 | 4 | EVNSFKAALSSLAKH | 2073 | 31 |
| HLA-DQA1*05:01/DQB1*02:01 | 2 | KNEVNSFKAALSSLA | 3341 | 32 |
| HLA-DQA1*01:01/DQB1*05:01 | 1 | VKNEVNSFKAALSSL | 3922 | 29 |

Table 4: Database-predicted HLA binding affinities for VKNEVNSFKAALSSLAKHG, the class II peptide used in this study.

Computed H2-I Binding Affinities for VKNEVNSFKAALSSLAKHG

| Allele | Start | 15-mer peptide | Median $pIC_{50}$nM | Consensus Score |
|---|---|---|---|---|
| **$H2\text{-}IA^{b}$** | 4 | EVNSFKAALSSLAKH | 138 | 1.4 |
| $H2\text{-}IA^{d}$ | 5 | VNSFKAALSSLAKHG | 1069 | 6.1 |
| $H2\text{-}IE^{d}$ | 5 | VNSFKAALSSLAKHG | 5797 | 35 |

Table 5: Database-predicted H2-I binding affinities for VKNEVNSFKAALSSLAKHG, the class II peptide used in this study.

```
QHO62884        QASSRSSSRSRNSXRNSTPGSSRGTSPARMAGNGGDAALALLLLDRLNQLESKMSGKGQQ  240
QIC50514        QASSRSSSRSRNSSRNSTPGSNRGTSPARMAGNGGDAALALLLLDRLNQLESKMSGKGQQ  240
QIC50515        QASSRSSSRSRNSSRNSTPGSNRGTSPARMAGNGGDAALALLLLDRLNQLESKMSGKGQQ  240
QHZ00406        QASSRSSSRSRNSLRNSTPGSSRGTSPARMAGNGGDAALALLLLDRLNQLESKMSGKGQQ  240
QHW06046        QASSRSSSRSRNSLRNSTPGSSRGTSPARMAGNGGDAALALLLLDRLNQLESKMSGKGQQ  240
QHW06056        QASSRSSSRSRNSLRNSTPGSSRGTSPARMAGNGGDAALALLLLDRLNQLESKMSGKGQQ  240
QIA98602        QASSRSSSRSRNSSRNSTPGSSRGTSPARMAGNGGDAALALLLLDRLNQLESKMSGKGQQ  240
QIA20052        QASSRSSSRSRNSSRNSTPGSSRGTSPARMAGNGGDAALALLLLDRLNQLESKMSGKGQQ  240
QIA98613        QASSRSSSRSRNSSRNSTPGSSRGTSPARMAGNGGDAALALLLLDRLNQLESKMSGKGQQ  240
QHZ87599        QASSRSSSRSRNSSRNSTPGSSRGTSPARMAGNGGDAALALLLLDRLNQLESKMSGKGQQ  240
QHZ87589        QASSRSSSRSRNSSRNSTPGSSRGTSPARMAGNGGDAALALLLLDRLNQLESKMSGKGQQ  240
QHQ82471        QASSRSSSRSRNSSRNSTPGSSRGTSPARMAGNGGDAALALLLLDRLNQLESKMSGKGQQ  240
QHR63278        QASSRSSSRSRNSSRNSTPGSSRGTSPARMAGNGGDAALALLLLDRLNQLESKMSGKGQQ  240
QHR63258        QASSRSSSRSRNSSRNSTPGSSRGTSPARMAGNGGDAALALLLLDRLNQLESKMSGKGQQ  240
QHZ00365        QASSRSSSRSRNSSRNSTPGSSRGTSPARMAGNGGDAALALLLLDRLNQLESKMSGKGQQ  240
BCA25661        QASSRSSSRSRNSSRNSTPGSSRGTSPARMAGNGGDAALALLLLDRLNQLESKMSGKGQQ  240
QHZ00386        QASSRSSSRSRNSSRNSTPGSSRGTSPARMAGNGGDAALALLLLDRLNQLESKMSGKGQQ  240
QHZ00396        QASSRSSSRSRNSSRNSTPGSSRGTSPARMAGNGGDAALALLLLDRLNQLESKMSGKGQQ  240
BCA25671        QASSRSSSRSRNSSRNSTPGSSRGTSPARMAGNGGDAALALLLLDRLNQLESKMSGKGQQ  240
BCA25681        QASSRSSSRSRNSSRNSTPGSSRGTSPARMAGNGGDAALALLLLDRLNQLESKMSGKGQQ  240
QHU36851        QASSRSSSRSRNSSRNSTPGSSRGTSPARMAGNGGDAALALLLLDRLNQLESKMSGKGQQ  240
QHO62110        QASSRSSSRSRNSSRNSTPGSSRGTSPARMAGNGGDAALALLLLDRLNQLESKMSGKGQQ  240
QHQ71970        QASSRSSSRSRNSSRNSTPGSSRGTSPARMAGNGGDAALALLLLDRLNQLESKMSGKGQQ  240
QHO62115        QASSRSSSRSRNSSRNSTPGSSRGTSPARMAGNGGDAALALLLLDRLNQLESKMSGKGQQ  240
QHO60601        QASSRSSSRSRNSSRNSTPGSSRGTSPARMAGNGGDAALALLLLDRLNQLESKMSGKGQQ  240
QHU36871        QASSRSSSRSRNSSRNSTPGSSRGTSPARMAGNGGDAALALLLLDRLNQLESKMSGKGQQ  240
QHW06066        QASSRSSSRSRNSSRNSTPGSSRGTSPARMAGNGGDAALALLLLDRLNQLESKMSGKGQQ  240
QHN73817        QASSRSSSRSRNSSRNSTPGSSRGTSPARMAGNGGDAALALLLLDRLNQLESKMSGKGQQ  240
QHU79181        QASSRSSSRSRNSSRNSTPGSSRGTSPARMAGNGGDAALALLLLDRLNQLESKMSGKGQQ  240
QHU79201        QASSRSSSRSRNSSRNSTPGSSRGTSPARMAGNGGDAALALLLLDRLNQLESKMSGKGQQ  240
QHU36831        QASSRSSSRSRNSSRNSTPGSSRGTSPARMAGNGGDAALALLLLDRLNQLESKMSGKGQQ  240
QHU36841        QASSRSSSRSRNSSRNSTPGSSRGTSPARMAGNGGDAALALLLLDRLNQLESKMSGKGQQ  240
QHU36861        QASSRSSSRSRNSSRNSTPGSSRGTSPARMAGNGGDAALALLLLDRLNQLESKMSGKGQQ  240
QHU79211        QASSRSSSRSRNSSRNSTPGSSRGTSPARMAGNGGDAALALLLLDRLNQLESKMSGKGQQ  240
QIC50512        QASSRSSSRSRNSSRNSTPGSSRGTSPARMAGNGGDAALALLLLDRLNQLESKMSGKGQQ  240
QHR63288        QASSRSSSRSRNSSRNSTPGSSRGTSPARMAGNGGDAALALLLLDRLNQLESKMSGKGQQ  240
QIC50508        QASSRSSSRSRNSSRNSTPGSSRGTSPARMAGNGGDAALALLLLDRLNQLESKMSGKGQQ  240
QIC50513        QASSRSSSRSRNSSRNSTPGSSRGTSPARMAGNGGDAALALLLLDRLNQLESKMSGKGQQ  240
QIC50516        QASSRSSSRSRNSSRNSTPGSSRGTSPARMAGNGGDAALALLLLDRLNQLESKMSGKGQQ  240
QIC50507        QASSRSSSRSRNSSRNSTPGSSRGTSPARMAGNGGDAALALLLLDRLNQLESKMSGKGQQ  240
QIC50509        QASSRSSSRSRNSSRNSTPGSSRGTSPARMAGNGGDAALALLLLDRLNQLESKMSGKGQQ  240
QIC50511        QASSRSSSRSRNSSRNSTPGSSRGTSPARMAGNGGDAALALLLLDRLNQLESKMSGKGQQ  240
QHD43423        QASSRSSSRSRNSSRNSTPGSSRGTSPARMAGNGGDAALALLLLDRLNQLESKMSGKGQQ  240
QIB84680        QASSRSSSRSRNSSRNSTPGSSRGTSPARMAGNGGDAALALLLLDRLNQLESKMSGKGQQ  240
QIC50510        QASSRSSSRSRNSSRNSTPGSSRGTSPARMAGNGGDAALALLLLDRLNQLESKMSGKGQQ  240
QHQ71980        QASSRSSSRSRNSSRNSTPGSSRGTSPARMAGNGGDAALALLLLDRLNQLESKMSGKGQQ  240
QHN73802        QASSRSSSRSRNSSRNSTPGSSRGTSPARMAGNGGDAALALLLLDRLNQLESKMSGKGQQ  240
QHR84456        QASSRSSSRSRNSSRNSTPGSSRGTSPARMAGNGGDAALALLLLDRLNQLESKMSGKGQQ  240
QHR63298        QASSRSSSRSRNSSRNSTPGSSRGTSPARMAGNGGDAALALLLLDRLNQLESKMSGKGQQ  240
YP_009724397    QASSRSSSRSRNSSRNSTPGSSRGTSPARMAGNGGDAALALLLLDRLNQLESKMSGKGQQ  240
QHR63268        QASSRSSSRSRNSSRNSTPGSSRGTSPARMAGNGGDAALALLLLDRLNQLESKMSGKGQQ  240
QIC53221        QASSRSSSRSRNSSRNSTPGSSRGTSPARMAGNGGDAALALLLLDRLNQLESKMSGKGQQ  240
QIC53211        QASSRSSSRSRNSSRNSTPGSSRGTSPARMAGNGGDAALALLLLDRLNQLESKMSGKGQQ  240
BCA25651        QASSRSSSRSRNSSRNSTPGSSRGTSPARMAGNGGDAALALLLLDRLNQLESKMSGKGQQ  240
                ************* *******.**************************************
```

Figure 7: Part 1 of 2. Sequences from 54 subjects with COVID-19 were found to have highly conserved nucleocapsid peptide sequences from positions 1-419 with the exception of three positions. At position 194, three individual sequences differ with non-conserved amino acid residues and one unknown amino acid. At position 202, a partially conserved amino acid variant is seen in two samples. At position 344, one non-conserved amino acid is present, however, this sample used a laboratory host cell line where only one of 4 replicates displayed this non-conserved amino acid substitution. These three mutation positions are colored according to the Clustal X color scheme.

```
QHO62884         WPQIAQFAPSASAFFGMSRIGMEVTPSGTWLTYTGAIKLDDKDPNFKDQVILLNKHIDAY  360
QIC50514         WPQIAQFAPSASAFFGMSRIGMEVTPSGTWLTYTGAIKLDDKDPNFKDQVILLNKHIDAY  360
QIC50515         WPQIAQFAPSASAFFGMSRIGMEVTPSGTWLTYTGAIKLDDKDPNFKDQVILLNKHIDAY  360
QHZ00406         WPQIAQFAPSASAFFGMSRIGMEVTPSGTWLTYTGAIKLDDKDPNFKDQVILLNKHIDAY  360
QHW06046         WPQIAQFAPSASAFFGMSRIGMEVTPSGTWLTYTGAIKLDDKDPNFKDQVILLNKHIDAY  360
QHW06056         WPQIAQFAPSASAFFGMSRIGMEVTPSGTWLTYTGAIKLDDKDPNFKDQVILLNKHIDAY  360
QIA98602         WPQIAQFAPSASAFFGMSRIGMEVTPSGTWLTYTGAIKLDDKDPNFKDQVILLNKHIDAY  360
QIA20052         WPQIAQFAPSASAFFGMSRIGMEVTPSGTWLTYTGAIKLDDKDPNFKDQVILLNKHIDAY  360
QIA98613         WPQIAQFAPSASAFFGMSRIGMEVTPSGTWLTYTGAIKLDDKDPNFKDQVILLNKHIDAY  360
QHZ87599         WPQIAQFAPSASAFFGMSRIGMEVTPSGTWLTYTGAIKLDDKDPNFKDQVILLNKHIDAY  360
QHZ87589         WPQIAQFAPSASAFFGMSRIGMEVTPSGTWLTYTGAIKLDDKDPNFKDQVILLNKHIDAY  360
QHQ82471         WPQIAQFAPSASAFFGMSRIGMEVTPSGTWLTYTGAIKLDDKDPNFKDQVILLNKHIDAY  360
QHR63278         WPQIAQFAPSASAFFGMSRIGMEVTPSGTWLTYTGAIKLDDKDPNFKDQVILLNKHIDAY  360
QHR63258         WPQIAQFAPSASAFFGMSRIGMEVTPSGTWLTYTGAIKLDDKDPNFKDQVILLNKHIDAY  360
QHZ00365         WPQIAQFAPSASAFFGMSRIGMEVTPSGTWLTYTGAIKLDDKDPNFKDQVILLNKHIDAY  360
BCA25661         WPQIAQFAPSASAFFGMSRIGMEVTPSGTWLTYTGAIKLDDKDPNFKDQVILLNKHIDAY  360
QHZ00386         WPQIAQFAPSASAFFGMSRIGMEVTPSGTWLTYTGAIKLDDKDPNFKDQVILLNKHIDAY  360
QHZ00396         WPQIAQFAPSASAFFGMSRIGMEVTPSGTWLTYTGAIKLDDKDPNFKDQVILLNKHIDAY  360
BCA25671         WPQIAQFAPSASAFFGMSRIGMEVTPSGTWLTYTGAIKLDDKDPNFKDQVILLNKHIDAY  360
BCA25681         WPQIAQFAPSASAFFGMSRIGMEVTPSGTWLTYTGAIKLDDKDPNFKDQVILLNKHIDAY  360
QHU36851         WPQIAQFAPSASAFFGMSRIGMEVTPSGTWLTYTGAIKLDDKDPNFKDQVILLNKHIDAY  360
QHO62110         WPQIAQFAPSASAFFGMSRIGMEVTPSGTWLTYTGAIKLDDKDPNFKDQVILLNKHIDAY  360
QHQ71970         WPQIAQFAPSASAFFGMSRIGMEVTPSGTWLTYTGAIKLDDKDPNFKDQVILLNKHIDAY  360
QHO62115         WPQIAQFAPSASAFFGMSRIGMEVTPSGTWLTYTGAIKLDDKDPNFKDQVILLNKHIDAY  360
QHO60601         WPQIAQFAPSASAFFGMSRIGMEVTPSGTWLTYTGAIKLDDKDPNFKDQVILLNKHIDAY  360
QHU36871         WPQIAQFAPSASAFFGMSRIGMEVTPSGTWLTYTGAIKLDDKDPNFKDQVILLNKHIDAY  360
QHW06066         WPQIAQFAPSASAFFGMSRIGMEVTPSGTWLTYTGAIKLDDKDPNFKDQVILLNKHIDAY  360
QHN73817         WPQIAQFAPSASAFFGMSRIGMEVTPSGTWLTYTGAIKLDDKDPNFKDQVILLNKHIDAY  360
QHU79181         WPQIAQFAPSASAFFGMSRIGMEVTPSGTWLTYTGAIKLDDKDPNFKDQVILLNKHIDAY  360
QHU79201         WPQIAQFAPSASAFFGMSRIGMEVTPSGTWLTYTGAIKLDDKDPNFKDQVILLNKHIDAY  360
QHU36831         WPQIAQFAPSASAFFGMSRIGMEVTPSGTWLTYTGAIKLDDKDPNFKDQVILLNKHIDAY  360
QHU36841         WPQIAQFAPSASAFFGMSRIGMEVTPSGTWLTYTGAIKLDDKDPNFKDQVILLNKHIDAY  360
QHU36861         WPQIAQFAPSASAFFGMSRIGMEVTPSGTWLTYTGAIKLDDKDPNFKDQVILLNKHIDAY  360
QHU79211         WPQIAQFAPSASAFFGMSRIGMEVTPSGTWLTYTGAIKLDDKDPNFKDQVILLNKHIDAY  360
QIC50512         WPQIAQFAPSASAFFGMSRIGMEVTPSGTWLTYTGAIKLDDKDPNFKDQVILLNKHIDAY  360
QHR63288         WPQIAQFAPSASAFFGMSRIGMEVTPSGTWLTYTGAIKLDDKDPNFKDQVILLNKHIDAY  360
QIC50508         WPQIAQFAPSASAFFGMSRIGMEVTPSGTWLTYTGAIKLDDKDPNFKDQVILLNKHIDAY  360
QIC50513         WPQIAQFAPSASAFFGMSRIGMEVTPSGTWLTYTGAIKLDDKDPNFKDQVILLNKHIDAY  360
QIC50516         WPQIAQFAPSASAFFGMSRIGMEVTPSGTWLTYTGAIKLDDKDPNFKDQVILLNKHIDAY  360
QIC50507         WPQIAQFAPSASAFFGMSRIGMEVTPSGTWLTYTGAIKLDDKDPNFKDQVILLNKHIDAY  360
QIC50509         WPQIAQFAPSASAFFGMSRIGMEVTPSGTWLTYTGAIKLDDKDPNFKDQVILLNKHIDAY  360
QIC50511         WPQIAQFAPSASAFFGMSRIGMEVTPSGTWLTYTGAIKLDDKDPNFKDQVILLNKHIDAY  360
QHD43423         WPQIAQFAPSASAFFGMSRIGMEVTPSGTWLTYTGAIKLDDKDPNFKDQVILLNKHIDAY  360
QIB84680         WPQIAQFAPSASAFFGMSRIGMEVTPSGTWLTYTGAIKLDDKDPNFKDQVILLNKHIDAY  360
QIC50510         WPQIAQFAPSASAFFGMSRIGMEVTPSGTWLTYTGAIKLDDKDPNFKDQVILLNKHIDAY  360
QHQ71980         WPQIAQFAPSASAFFGMSRIGMEVTPSGTWLTYTGAIKLDDKDPNFKDQVILLNKHIDAY  360
QHN73802         WPQIAQFAPSASAFFGMSRIGMEVTPSGTWLTYTGAIKLDDKDPNFKDQVILLNKHIDAY  360
QHR84456         WPQIAQFAPSASAFFGMSRIGMEVTPSGTWLTYTGAIKLDDKDPNFKDQVILLNKHIDAY  360
QHR63298         WPQIAQFAPSASAFFGMSRIGMEVTPSGTWLTYTGAIKLDDKDPNFKDQVILLNKHIDAY  360
YP_009724397     WPQIAQFAPSASAFFGMSRIGMEVTPSGTWLTYTGAIKLDDKDPNFKDQVILLNKHIDAY  360
QHR63268         WPQIAQFAPSASAFFGMSRIGMEVTPSGTWLTYTGAIKLDDKDPNFKDQVILLNKHIDAY  360
QIC53221         WPQIAQFAPSASAFFGMSRIGMEVTPSGTWLTYTGAIKLDDKDPNFKDQVILLNKHIDAY  360
QIC53211         WPQIAQFAPSASAFFGMSRIGMEVTPSGTWLTYTGAIKLDDKDPNFKDQVILLNKHIDAY  360
BCA25651         WPQIAQFAPSASAFFGMSRIGMEVTPSGTWLTYTGAIKLDDKDSNFKDQVILLNKHIDAY  360
                 ******************************************* ****************
```

Figure 8: Part 2 of 2. Sequences from 54 subjects with COVID-19 were found to have highly conserved nucleocapsid peptide sequences from positions 1-419 with the exception of three positions. At position 194, three individual sequences differ with non-conserved amino acid residues and one unknown amino acid. At position 202, a partially conserved amino acid variant is seen in two samples. At position 344, one non-conserved amino acid is present, however, this sample used a laboratory host cell line where only one of 4 replicates displayed this non-conserved amino acid substitution. These three mutation positions are colored according to the Clustal X color scheme.

COVID-19 Nucleocapsid Peptides with Associated Predicted HLA Restricted Binding Affinities (1/4)

| Peptide | Start Position | Allele | NetMHC 4.0 $pIC_{50}nM$ | NetMHCpan 4.0 $pIC_{50}nM$ | SARS Same? |
|---|---|---|---|---|---|
| LSPRWYFYY | 104 | HLA-A*01:01 | 48.64 | 76.9 | YES |
| LLLDRLNQL | 222 | HLA-A*02:01 | 14.81 | 11.3 | YES |
| GMSRIGMEV | 316 | HLA-A*02:01 | 50.61 | 48.1 | YES |
| KTFPPTEPK | 361 | HLA-A*03:01 | 20.8 | 18.8 | YES |
| KSAAEASKK | 249 | HLA-A*03:01 | 116.22 | 139.4 | YES |
| LIRQGTDYK | 291 | HLA-A*03:01 | 274.69 | 137.5 | YES |
| ASAFFGMSR | 311 | HLA-A*03:01 | 292.41 | 285.3 | YES |
| QLESKMSGK | 229 | HLA-A*03:01 | 322.41 | 751 | NO |
| FTALTQHGK | 53 | HLA-A*03:01 | 788.84 | 345.5 | YES |
| KTFPPTEPK | 361 | HLA-A*11:01 | 6.28 | 7.7 | YES |
| ASAFFGMSR | 311 | HLA-A*11:01 | 14.4 | 15.3 | YES |
| FTALTQHGK | 53 | HLA-A*11:01 | 127.28 | 44.9 | YES |
| KSAAEASKK | 249 | HLA-A*11:01 | 76.73 | 62.2 | YES |
| AGLPYGANK | 119 | HLA-A*11:01 | 240.23 | 157.5 | NO |
| LIRQGTDYK | 291 | HLA-A*11:01 | 984.82 | 160.6 | YES |
| LSPRWYFYY | 104 | HLA-A*11:01 | 253.34 | 492.8 | YES |
| TQALPQRQK | 379 | HLA-A*11:01 | 740.66 | 415.1 | NO |
| QQQGQTVTK | 240 | HLA-A*11:01 | 428.26 | 470.3 | YES |
| KHIDAYKTF | 355 | HLA-A*23:01 | 134.12 | 778.7 | YES |
| YYRRATRRI | 86 | HLA-A*23:01 | 151.38 | 366.6 | NO |
| TWLTYTGAI | 329 | HLA-A*23:01 | 24164.38 | 282.1 | NO |
| KHWPQIAQF | 299 | HLA-A*23:01 | 317.71 | 313.7 | YES |
| KAYNVTQAF | 266 | HLA-A*23:01 | 341.14 | 602.3 | NO |
| YYRRATRRI | 86 | HLA-A*24:02 | 74.89 | 322 | NO |

Table 6: This set of 53 unique peptides (part 1 of 4) achieves > 95% world-wide population coverage. The starting position is within the nucleocapsid. Peptides chosen with binding affinity predictions less than 500nm via NetMHC 4.0 or NetMHCpan 4.0. Peptide sequences colored in red have literature references as known *in-vitro* binders to the predicted allele match (see text).

### COVID-19 Nucleocapsid Peptides with Associated Predicted HLA Restricted Binding Affinities (2/4)

| Peptide | Start Position | Allele | NetMHC 4.0 $pIC_{50}nM$ | NetMHCpan 4.0 $pIC_{50}nM$ | SARS Same? |
|---|---|---|---|---|---|
| FAPSASAFF | 307 | HLA-A*24:02 | 422.31 | 847.7 | YES |
| NTASWFTAL | 48 | HLA-A*26:01 | 1113.04 | 122.6 | YES |
| ELIRQGTDY | 290 | HLA-A*26:01 | 652.8 | 327.8 | NO |
| FAPSASAFF | 307 | HLA-A*26:01 | 349.57 | 606.6 | YES |
| IGYYRRATR | 84 | HLA-A*33:03 | N/A | 57.8 | YES |
| NVTQAFGRR | 269 | HLA-A*33:03 | N/A | 62.5 | YES |
| ASAFFGMSR | 311 | HLA-A*33:03 | N/A | 149.3 | YES |
| QASSRSSSR | 181 | HLA-A*33:03 | N/A | 163.9 | YES |
| YNVTQAFGR | 268 | HLA-A*33:03 | N/A | 189.1 | YES |
| GYYRRATRR | 85 | HLA-A*33:03 | N/A | 359.4 | YES |
| SSRSSSRSR | 183 | HLA-A*33:03 | N/A | 395.3 | YES |
| FPRGQGVPI | 66 | HLA-B*07:02 | 3.82 | 4.7 | YES |
| KPRQKRTAT | 257 | HLA-B*07:02 | 4.42 | 18.8 | YES |
| SPRWYFYYL | 105 | HLA-B*07:02 | 6.32 | 15.3 | YES |
| RIRGGDGKM | 93 | HLA-B*07:02 | 149.86 | 173 | NO |
| NPANNAAIV | 150 | HLA-B*07:02 | 184.8 | 569.3 | NO |
| LPNNTASWF | 45 | HLA-B*07:02 | 244.3 | 334 | YES |
| SPRWYFYYL | 105 | HLA-B*08:01 | 13.77 | 42.1 | YES |
| LLLDRLNQL | 222 | HLA-B*08:01 | 125.72 | 136.8 | YES |
| FPRGQGVPI | 66 | HLA-B*08:01 | 245.35 | 368.3 | YES |
| KPRQKRTAT | 257 | HLA-B*08:01 | 364.72 | 432.6 | YES |
| KAYNVTQAF | 266 | HLA-B*15:01 | 40.35 | 19 | NO |
| LLNKHIDAY | 352 | HLA-B*15:01 | 33.04 | 32.5 | YES |

Table 7: This set of 53 unique peptides (part 2 of 4) achieves > 95% world-wide population coverage. The starting position is within the nucleocapsid. Peptides chosen with binding affinity predictions less than 500nm via NetMHC 4.0 or NetMHCpan 4.0. Peptide sequences colored in red have literature references as known *in-vitro* binders to the predicted allele match (see text).

COVID-19 Nucleocapsid Peptides with Associated Predicted HLA Restricted Binding Affinities (3/4)

| Peptide | Start Position | Allele | NetMHC 4.0 $pIC_{50}nM$ | NetMHCpan 4.0 $pIC_{50}nM$ | SARS Same? |
|---|---|---|---|---|---|
| LQLPQGTTL | 159 | HLA-B*15:01 | 105.55 | 229.8 | YES |
| FAPSASAFF | 307 | HLA-B*15:01 | 213.11 | 281.9 | YES |
| FSKQLQQSM | 403 | HLA-B*15:01 | 219.07 | 286 | NO |
| RLNQLESKM | 226 | HLA-B*15:01 | 1496.11 | 490.3 | NO |
| QFAPSASAF | 306 | HLA-B*15:01 | 493.85 | 700.3 | YES |
| RRIRGGDGK | 92 | HLA-B*27:05 | 65.94 | 72.5 | NO |
| RRATRRIRG | 88 | HLA-B*27:05 | 253.64 | 787.8 | NO |
| QRNAPRITF | 9 | HLA-B*27:05 | 560.56 | 262.1 | NO |
| YRRATRRIR | 87 | HLA-B*27:05 | 415.31 | 597.7 | NO |
| NTASWFTAL | 48 | HLA-B*39:01 | 47.87 | 353.3 | YES |
| KKADETQAL | 374 | HLA-B*39:01 | 137.43 | 926.4 | NO |
| LQLPQGTTL | 159 | HLA-B*39:01 | 238.19 | 228.7 | YES |
| TRNPANNAA | 148 | HLA-B*39:01 | 406.62 | 818.3 | NO |
| MEVTPSGTW | 322 | HLA-B*44:02 | 11.48 | 14.2 | YES |
| LPNNTASWF | 45 | HLA-B*53:01 | 19.03 | 25.7 | YES |
| TPSGTWLTY | 325 | HLA-B*53:01 | 26.99 | 79 | YES |
| LPAADLDDF | 395 | HLA-B*53:01 | 193.75 | 74.8 | NO |
| FAPSASAFF | 307 | HLA-B*53:01 | 1164.6 | 317.4 | YES |
| GANKDGIIW | 124 | HLA-B*53:01 | 320.56 | 1015.8 | NO |
| KAYNVTQAF | 266 | HLA-B*58:01 | 12.51 | 17.7 | NO |
| GANKDGIIW | 124 | HLA-B*58:01 | 158.07 | 35.3 | NO |
| KMKDLSPRW | 100 | HLA-B*58:01 | 83.99 | 99.2 | NO |
| LSPRWYFYY | 104 | HLA-B*58:01 | 359.42 | 430.6 | YES |
| KAYNVTQAF | 266 | HLA-C*03:04 | N/A | 12.7 | NO |

Table 8: This set of 53 unique peptides (part 3 of 4) achieves $> 95\%$ world-wide population coverage. The starting position is within the nucleocapsid. Peptides chosen with binding affinity predictions less than 500nm via NetMHC 4.0 or NetMHCpan 4.0. Peptide sequences colored in red have literature references as known *in-vitro* binders to the predicted allele match (see text).

COVID-19 nucleocapsid Peptides with Associated Predicted HLA Restricted Binding Affinities (4/4)

| Peptide | Start Position | Allele | NetMHC 4.0 $pIC_{50}nM$ | NetMHCpan 4.0 $pIC_{50}nM$ | SARS Same? |
|---|---|---|---|---|---|
| FAPSASAFF | 307 | HLA-C*03:04 | N/A | 41.4 | YES |
| LTYTGAIKL | 331 | HLA-C*03:04 | N/A | 44.8 | NO |
| NTASWFTAL | 48 | HLA-C*03:04 | N/A | 58.8 | YES |
| SAFFGMSRI | 312 | HLA-C*03:04 | N/A | 68 | YES |
| LQLPQGTTL | 159 | HLA-C*03:04 | N/A | 99.5 | YES |
| FSKQLQQSM | 403 | HLA-C*03:04 | N/A | 149.9 | NO |
| FPRGQGVPI | 66 | HLA-C*03:04 | N/A | 434.9 | YES |
| YRRATRRIR | 87 | HLA-C*07:01 | 112.27 | 8786.2 | NO |
| QRNAPRITF | 9 | HLA-C*07:01 | 1337.36 | 198.9 | NO |
| YYRRATRRI | 86 | HLA-C*07:01 | 254.32 | 957.2 | NO |
| LKFPRGQGV | 64 | HLA-C*07:01 | 446.18 | 1633.3 | NO |
| QRNAPRITF | 9 | HLA-C*07:02 | 261.17 | 237.8 | NO |
| YYRRATRRI | 86 | HLA-C*07:02 | 6229.2 | 242.2 | NO |
| FAPSASAFF | 307 | HLA-C*07:02 | 16893.5 | 347.4 | YES |
| KHWPQIAQF | 299 | HLA-C*07:02 | 430.68 | 971.3 | YES |
| NFKDQVILL | 345 | HLA-C*07:02 | 29905.43 | 462.1 | NO |
| KAYNVTQAF | 266 | HLA-C*07:02 | 668.01 | 477 | NO |
| FAPSASAFF | 307 | HLA-C*08:01 | N/A | 280.1 | YES |
| KAYNVTQAF | 266 | HLA-C*08:01 | N/A | 412.2 | NO |

Table 9: This set of 53 unique peptides (part 4 of 4) achieves $> 95\%$ world-wide population coverage. The starting position is within the nucleocapsid. Peptides chosen with binding affinity predictions less than 500nm via NetMHC 4.0 or NetMHCpan 4.0. Peptide sequences colored in red have literature references as known *in-vitro* binders to the predicted allele match (see text).

### Projected World-Wide Population Coverage for a COVID-19 Peptide Vaccine Targeting 9mer Peptides on Nucleocapsid Proteins

| Minimum Epitope Matches / Allele / Person | % Projected Coverage | Cumulative % Population Coverage |
|---|---|---|
| 1 | 18.14 | 97.27 |
| 2 | 35.05 | 79.13 |
| 3 | 29.7 | 44.08 |
| 4 | 12.02 | 14.37 |
| 5 | 2.2 | 2.35 |
| 6 | 0.15 | 0.15 |

Table 10: Data showing projected HLA world-wide population coverage for a COVID-19 vaccine using the 16 epitopes listed in Tables 6 through 9. If we assume a least one HLA match per peptide capable of producing a clinically relevant immune response in a person, we can achieve 97.27% global population coverage with a 16 class I peptide CTL vaccine.

### Projected China-Specific Population Coverage for a COVID-19 Peptide Vaccine Targeting 9mer Peptides on Nucleocapsid Proteins

| Minimum Epitope Matches / Allele / Person | % Projected Coverage | Cumulative % Population Coverage |
|---|---|---|
| 1 | 26.02 | 94.39 |
| 2 | 38.01 | 68.37 |
| 3 | 23.14 | 30.36 |
| 4 | 6.42 | 7.22 |
| 5 | 0.77 | 0.8 |
| 6 | 0.03 | 0.03 |

Table 11: If we take the assumptions made in the global projected population coverage Table 10 now assuming a China-specific HLA distribution, we still can achieve 94.39% population coverage

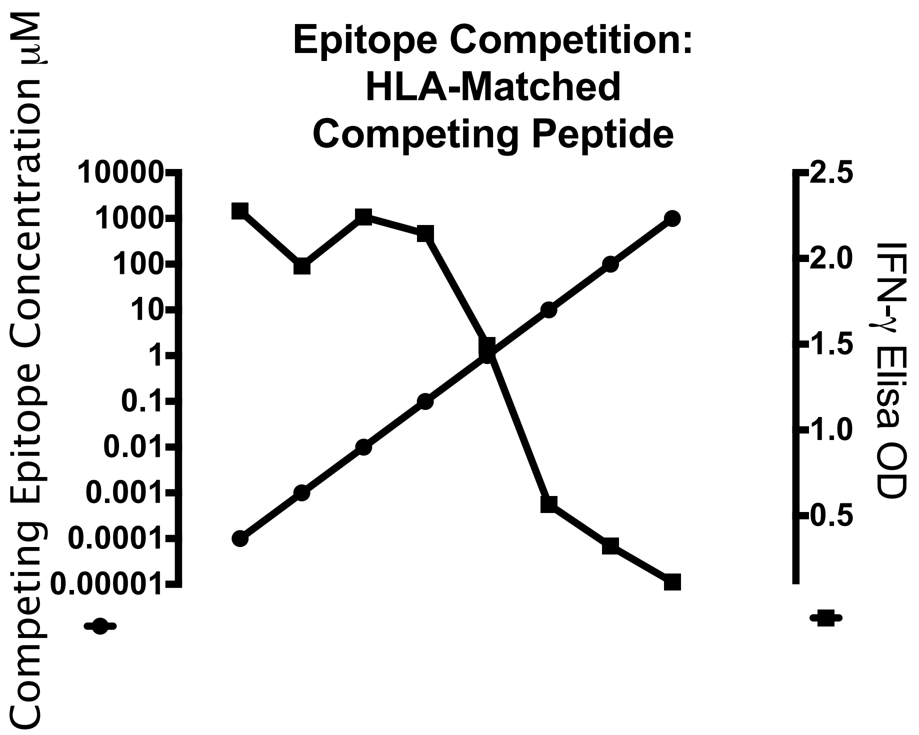

Figure 9: Simultaneous *in-vitro* incubation of two HLA matched epitopes to human PBMCs from an indivdual vaccinated against tetanus toxin. $IFN - \gamma$ release in response to a Classs I epitope from tetanus virus is inhibited by increasing concentrations of Class I epitope from HIV in a patient not exposed to the HIV virus.

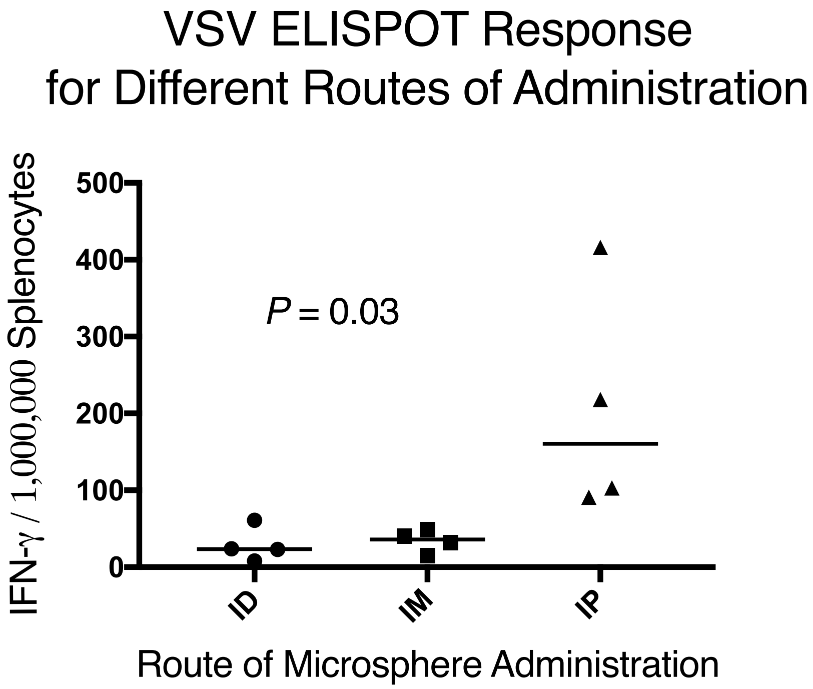

Figure 10: Differential ELISPOT response to intradermal tail (ID) intramuscular (IM) and intraperitoneal administration of 2mg of adjuvanted microspheres loaded with VSV Class I epitope RGYVYQGL.

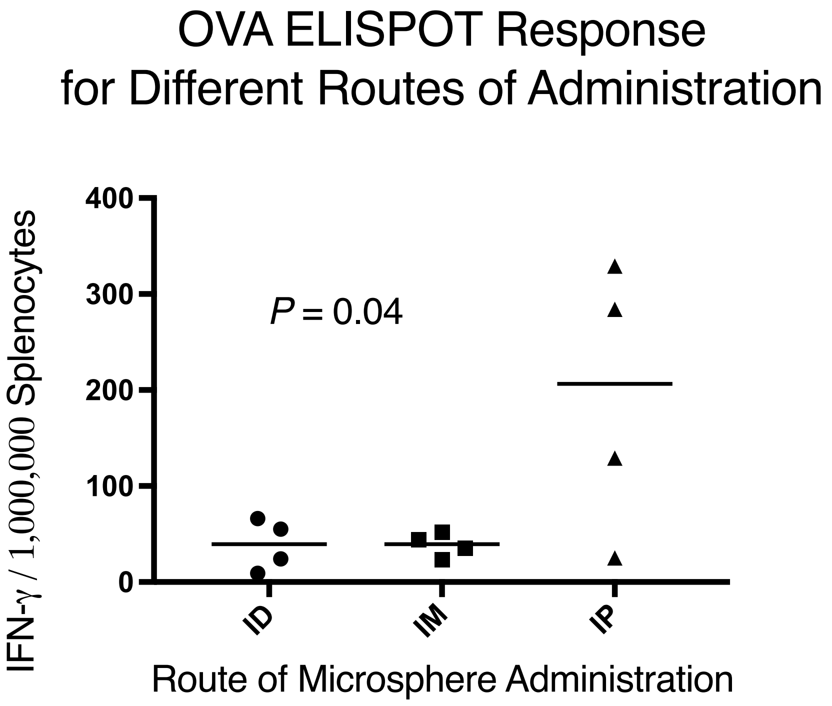

Figure 11: Differential ELISPOT response to intradermal tail (ID) intramuscular (IM) and intraperitoneal administration of 2mg of adjuvanted microspheres loaded with OVA Class I epitope RGYVVYQGL.

| Adjuvanted Microsphere Loaded Component | Amount |
|---|---|
| Peptide | 0.1% w/w |
| CpG | 0.025% w/w |
| Mannose | 0.01% w/w |

Table 12: Compositon of 11$\mu$M PLGA (Resomer 502H) adjuvanted microspheres used for the study.

| Injectate Component | Amount |
|---|---|
| Polysorbate 20 | 0.01% (v/v) |
| MPLA | 50$\mu$g/ml |

Table 13: Compositon of PBS injectate used for the study.

| Peptide Sequence | Description |
|---|---|
| SIINFEKL | OVA class I |
| ISQAVHAAHAEINEAGR | OVA class II |
| RGYVYQGL | VSV class I |
| SSKAGVFEHPHIGDASSGL | VSV class II |

Table 14: Class I and class II peptides used in the study. Any single microsphere used to inoculate mice for this study contained only one of the epitopes in this table.
